# Multiplex Gene Editing Suppresses Random Integration of Hepatitis B Virus DNA in Chronically Infected Liver

**DOI:** 10.1101/2025.05.06.652508

**Authors:** Samuel Slattery, Wenwen Huo, Amena Arif, Jennifer Gordon, Ryo Takeuchi

## Abstract

Gene editing technologies have opened the possibility of directly targeting viral DNA in therapeutic applications. In chronically infected hepatocytes with hepatitis B virus (HBV), covalently closed circular DNA (cccDNA) serves as the master template for viral transcripts and gene products. In the present study, we evaluated the outcomes of anti-HBV multiplex gene editing with the CRISPR-Cas9 endonuclease from Staphylococcus aureus (SaCas9) using primary human hepatocytes (PHHs) and HBV mouse models. Nonviral delivery of SaCas9-encoding mRNA and a pair of HBV-targeting guide RNAs (gRNAs) substantially reduced viral biomarkers and intrahepatic HBV DNA copies in vitro and in vivo, suggesting that fragmentation of HBV DNA primarily leads to its degradation. Hybridization capture sequencing analyses indicated that small insertions and deletions (indels) and structural variants including excisions and inversions of the viral sequences were accumulated in the residual HBV DNA. These assays also demonstrated that transient expression of the HBV-targeting SaCas9 significantly suppressed random integration of HBV DNA, while this therapeutic approach was unlikely to affect chromosomal translocations involving viral copies. Taken together, our results suggest that anti-HBV multiplex gene editing eliminates viral DNA from chronically infected hepatocytes, potentially reducing the risk of hepatocarcinogenesis associated with HBV DNA integration.

## Introduction

HBV is a hepatotropic DNA virus that carries 3.2-kb of the partially double-stranded, relaxed circular DNA (rcDNA) in the capsid proteins covered by the lipid envelop layer containing the three isoforms of hepatitis B surface antigen (HBsAg) ^1^. HBV enters into liver hepatocytes by binding to the sodium taurocholate cotransporting polypeptide (NTCP) via the unique, extended N-terminal peptide carried by the largest HBsAg isoform. The rcDNA coated by the capsid proteins is transported through the cytolytic trafficking machinery into the nuclei, where it is converted to cccDNA by the host proteins involved in DNA replication and repair. This form of HBV DNA persists in the nuclei and serves as a template for 5 types of mRNA encoding a total of 7 viral gene products. Among these viral transcripts, pregenomic RNA (pgRNA) is used as the template of reverse transcription by the viral polymerase during encapsidation.

HBV infection can lead to chronic hepatitis B (CHB), which is estimated to affect approximately 250 million people in the world ^2,3^ Persistence of HBV significantly increases the risk of developing liver diseases including cirrhosis, liver failure and hepatocellular carcinoma (HCC). HBV infection is the leading cause of HCC and accounts for >50 % of its cases worldwide. HBV DNA integration is one of the mechanisms that potentially contribute to hepatocarcinogenesis, and the integrated DNA (intDNA) copies have been found near HCC-associated genes including TERT and CCNE1 encoding telomerase reverse transcriptase and cyclin E1, respectively ^4,5^. The viral DNA can be inserted into genomic DNA early in the course of its infection, and the frequency of this event has been estimated to range between 0.01 % and 0.1 % under the experimental conditions tested previously ^6^. The copy number of integrated HBV DNA has also been observed to increase over the course of chronic infection ^7^. The double-stranded linear DNA (dslDNA) is a by-product created during viral replication and a preferential substrate used for integration in the nuclei ^8,9^. Nucleos(t)ide analogues (NAs), which are the standard of care for CHB, target viral reverse transcription and reduce the viral load by several orders of magnitude in patients with CHB ^10,11^. While this oral medication significantly prevents liver disease progression, it leads to functional cure at a very low rate. Functional cure is defined as sustained HBsAg loss and HBV DNA less than the lower limit of quantitation (LLOQ) 24 weeks off-treatment ^12,13^ and has been set as the preferred primary end point of clinical trials for novel treatments for CHB.

We previously sought to eliminate human immunodeficiency virus (HIV) and herpes simplex virus type-1 (HSV-1) by multiplex gene editing with CRISPR-Cas9 and demonstrated that systemic delivery of AAV vectors carrying the SaCas9 gene and a pair of gRNAs blocked viral rebound and reactivation in preclinical models ^14,15^. We further examined the HIV-targeting EBT-101 therapy in the clinic and confirmed its safety and tolerability (NCT05144386). These results suggest that it is feasible to treat chronic infection by directly removing viral DNA with CRISPR-Cas9. A similar therapeutic approach potentially offers a curative option for other infectious diseases including CHB. Systemic delivery of lipid nanoparticles (LNPs) encapsulating gene editing payloads has been demonstrated to introduce targeted gene disruption in hepatocytes of liver ^16,17^. Earlier studies have demonstrated that CRISPR-Cas9 can access intracellular HBV DNA to suppress viral transcription and protein expression ^18–23^, and two gene editing-based therapies for CHB have recently advanced to clinical trials (NCT06680232; NCT06671093). While anti-HBV gene editing therapy may benefit the lives of patients with CHB, its potential safety concerns include an increased frequency of HBV integration following linearization of episomal HBV DNA copies and unintended structural variants such as chromosomal translocations mediated by cleavage of intDNA. To investigate to what extent nuclease-mediated HBV DNA editing alters chromosomal DNA, we first selected a pair of gRNAs designed to target well-conserved sites of the viral genome and examined the anti-viral activity of SaCas9 complexed with each of the paired gRNAs in HBV-infected cells and in two CHB mouse models. We then analyzed DNA edits introduced by our therapeutic method in primary human hepatocytes and mouse tissue samples by deep sequencing. We found that our multiplexed HBV gene editing therapy greatly reduced viral antigen secretion, replication and random HBV DNA integration with little increased risk of chromosomal translocations involving the viral DNA. These results provided valuable insights into not only the molecular mechanisms of anti-HBV multiplex gene editing but also its impact on chromosomal integrity.

## Results

### Identifying a pair of guide RNAs that direct CRISPR/Cas9 to conserved sites of the HBV genome

HBV genomic sequences are highly diverse due to the high error rate during reverse transcription catalyzed by the viral polymerase that lacks proof-reading activity. Viral sequences that differ by >7.5 % are considered as separate genotypes, which show geographically different distributions ^24^. By in-silico analysis of viral sequences curated from HBVdb ^25^, we identified a pair of sites that could be targeted with SaCas9. One of these sites (SaCas9 gRNA 1 target site) is positioned in the region encoding the viral polymerase and major surface antigen, while the other (SaCas9 gRNA 2 target site) is in the core antigen gene. Both were well conserved without nucleotide substitutions and gaps in approximately 90 % or more of viral sequences annotated to each of 8 genotypes (A-H) in the database (Supplementary Fig. 1A). We next searched for homologous sequences on the human reference genome and found that only a few sites differed from these viral target sequences at 3 or fewer nucleotide positions within the 21-nt protospacer (Supplementary Table 1).

To assay on-target editing activity of SaCas9 with the selected pair of gRNAs, we generated an HBV reporter cell line carrying a synthetic reporter construct that contains the two HBV target sites separated by about 400 base pairs in the AAVS1 locus (Supplementary Fig. 1B). Transfection with SaCas9-encoding mRNA and the paired gRNAs introduced excision of the intervening region between the two target sites at 60 % of the chromosomally integrated reporter construct in the cell line (Supplementary Fig. 1C). We also sequenced the unexcised target sites and found that indels were formed at 45 % and 30 % of the PCR-amplified sites targeted by SaCas9 gRNA 1 and SaCas9 gRNA 2, respectively (Supplementary Fig. 1D). Since these target sites are located in the open reading frames of viral genes, both excisions and indels are predicted to disrupt viral protein expression.

To investigate specificity of gene editing with the selected gRNAs, we first nominated human genomic sites for off-target analysis by GUIDE-seq ^26,27^. We transfected the HBV reporter cell line with SaCas9-encoding mRNA, either of SaCas9 gRNA 1 or SaCas9 gRNA 2 and a double-stranded oligonucleotide (dsODN) by electroporation, then identified dsODN insertion sites on the human genome by next-generation sequencing. As expected, most sequence reads containing dsODN were mapped on the HBV on-target sites of the chromosomally integrated construct and only a small number of dsODNs were detected near human genomic sites that shared relatively high sequence identity with the intended target sites (Supplementary Figs. 2A,B). To determine if SaCas9 complexed with each of the paired gRNAs edited the sites nominated by GUIDE-seq, we quantified indels at 3-4 sites in the same reporter cell line transfected with SaCas9-encoding mRNA and one of the paired gRNAs using targeted amplicon sequencing. These sites were selected based on read count from the GUIDE-seq assays and the number of base pair substitutions and gaps in the 9-nucleotide PAM-proximal region, which were predicted to substantially suppress the SaCas9 endonuclease activity ^28^. Indels were found at the majority of the integrated on-target sites derived from the HBV genomic sequence in the edited cells compared to <0.1 % in the control samples, confirming robust editing at the intended target sites by our anti-HBV gene editing nuclease under the conditions tested (Supplementary Figs. 2C,D). In contrast, very little change in the frequency of indels was observed at any of the nominated sites between the treated and control samples. These results suggest that the selected gRNAs specifically direct SaCas9 to the viral target sequences.

### Multiplex gene editing-mediated suppression of HBV replication in HBV-infected primary human hepatocytes

To examine if our anti-HBV editing approach inhibited viral replication in primary human hepatocytes (PHHs), we transfected HBV-infected PHHs with SaCas9-encoding mRNA and the pair of HBV-targeting gRNAs at 4 days post-infection and analyzed viral antigens secreted to the culture medium and intrahepatic HBV DNA at 8 days and 11 days post-infection (Day 9 and Day 12) as shown in Fig. 1A. Chemiluminescent immunoassays showed that concentrations of HBsAg and HBeAg in the harvested culture media were lowered in the edited samples by 79 % and 75 %, respectively on Day 9 compared to samples transfected with a control gRNA (Figs 1B,C). We observed further reduction of HBsAg and HBeAg from the baseline in edited samples on Day 12. In addition, intrahepatic HBV DNA copies were reduced by 58 % and 71 % in cells transfected with the HBV-targeting gRNAs on Day 9 and Day 12, respectively (Fig. 1D). These results suggest that cleavage of viral DNA leads to its degradation and suppression of viral gene expression in hepatocytes.

**Figure 1.**
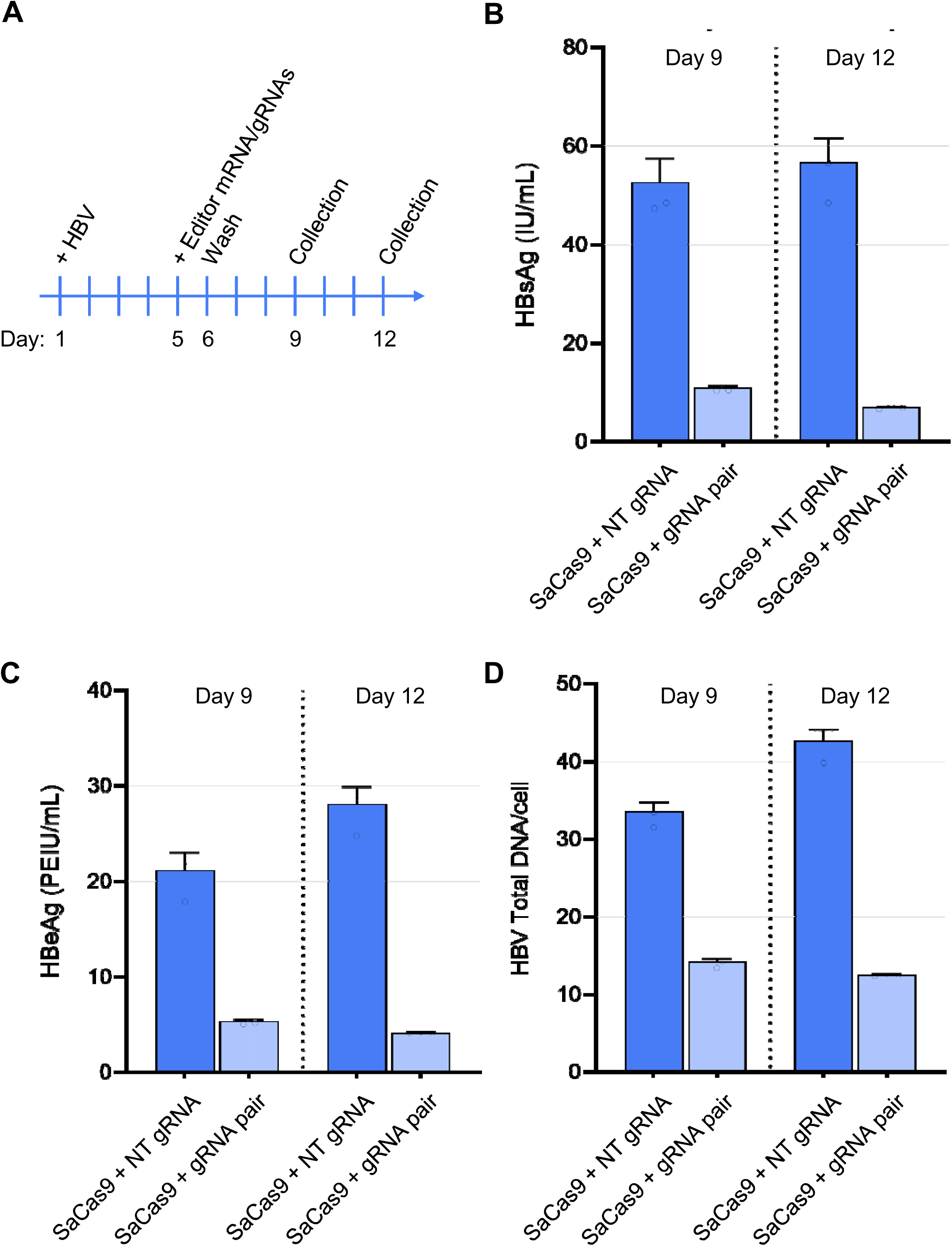
Anti-HBV multiple gene editing in HBV-infected PHHs. (A) A schematic of the experimental procedures. Primary hepatocytes were infected with HBV one day after cell seeding (Day 1), and transfection with the SaCas9-encoding mRNA and paired gRNAs was performed on Day 5. The culture media were exchanged on Days 6, 9 and 12. (B,C) Concentrations of HBsAg (B) and HBeAg (C) were quantified by chemiluminescent immunoassays. The control cultures were transfected with the non-targeting (NT) gRNA instead of a pair of the HBV-targeting gRNAs (gRNA pair). (D) Intrahepatic copies of total HBV DNA were quantified by digital PCR with DNA samples extracted from cells harvested on Day 9 or Day 12.

To characterize sequence alterations on the residual HBV DNA recovered from the edited cells, we performed hybridization capture sequencing with HBV-targeting capture probes. The sequencing depth was relatively even across the HBV genome, and its relative coverage to the internal control, human GAPDH, was lowered in the edited samples, as expected (Supplementary Fig. 3). In the viral sequences that escaped clearance, we detected indels at 1.8 and 1.1 % of the HBV on-target sites for SaCas9 gRNA 1 and SaCas9 gRNA 2, respectively (Fig. 2A). In addition, we observed structural variants that might be the result of excision and inversion of the viral sequences between the two target sites (Figs. 2B,C). These findings indicate that our HBV-targeting endonuclease can generate double strand breaks at the intended target sites of HBV DNA that persists in human hepatocytes. The majority of the cleaved DNA products may be eliminated prior to end-joining by the endogenous DNA repair machinery.

**Figure 2.**
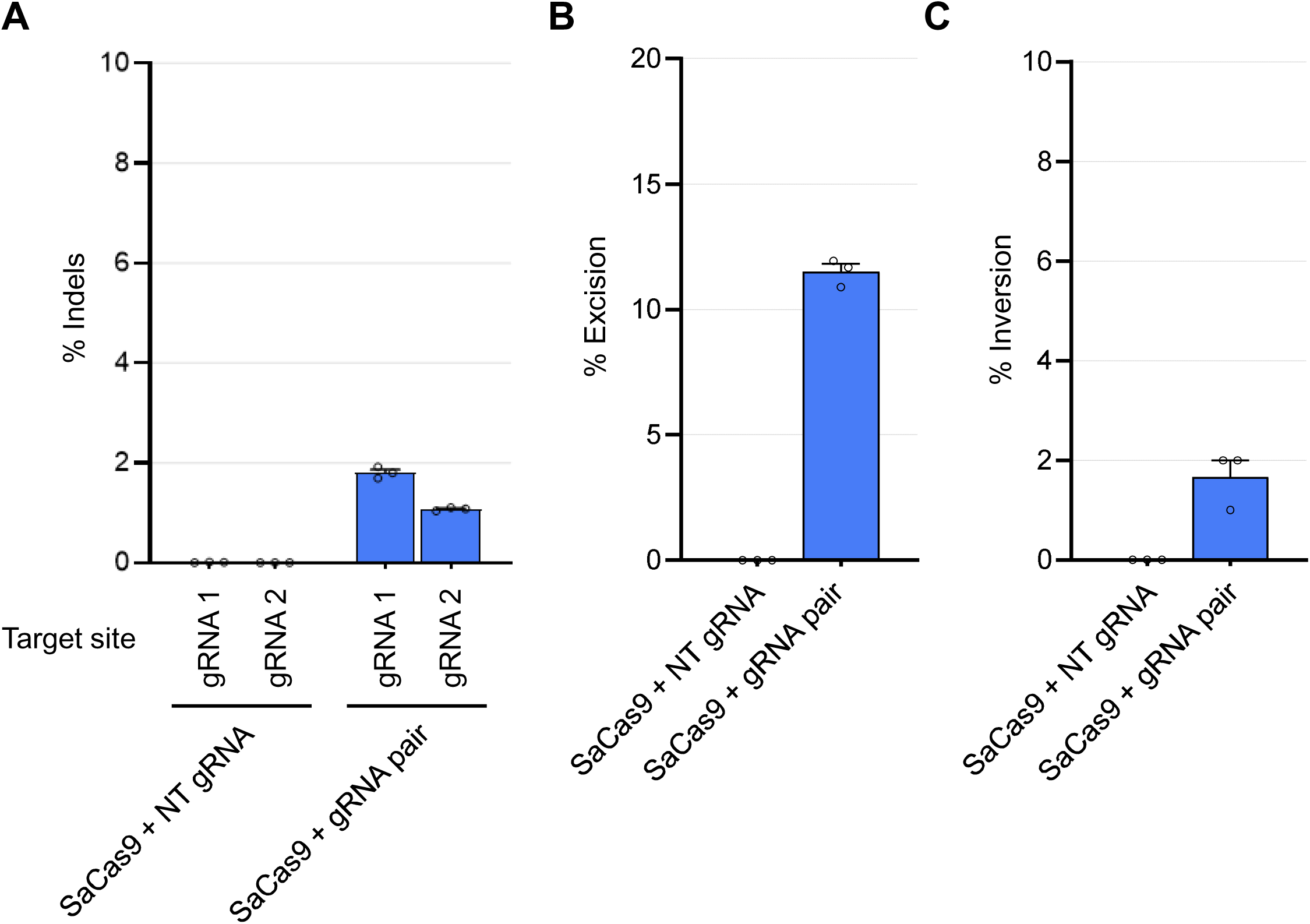
Sequence analysis of HBV DNA copies recovered from HBV-infected PHHs by short-read hybridization capture sequencing. (A) Frequency of indels were calculated by sequence reads containing indels at each on-target site by those carrying its flanking regions. (B,C) Sequence reads representing excisions (B) or inversions (C) of sequences between the two intended target sites were identified, and the percentage of such editing events were calculated using the number of total reads containing sequences adjacent to either of the intended cut sites.

HBV is an oncogenic virus that potentially induces chromosomal abnormalities through random DNA integration ^29^. A small number of sequence reads derived from these genetic events were found in both control and edited samples collected only 11 days post HBV infection. As described previously ^5,8,30^, the break points of the chromosomal insertion sites were distributed across the HBV genome and increased around Direct Repeat 1 (DR1) positioned upstream of the viral core antigen gene (nucleotide positions 1800-1860) (Fig. 3A). These sites were proximal to the predicted termini of dslDNA, which has been thought to serve as the primary source of randomly integrated HBV DNA. The relative number of HBV-chromosome junctions to the average sequence coverage of the internal control gene (human GAPDH) was reduced by editing with a pair of the HBV-targeting gRNAs (Fig. 3B), indicating that fragmentation of HBV DNA didn’t facilitate HBV DNA chromosomal insertion. This may be because our anti-HBV SaCas9 limited viral replication and dslDNA that was transported into the nuclei. A slightly increased number of break points were detected in the vicinity of the two intended target sites compared to the surrounding areas of the HBV genome in the edited samples. In cases where HBV DNA copies are inserted into chromosomes following fragmentation at the on-target site positioned on the HBsAg-encoding region, this gene product is unlikely to be expressed from the integrated sequences. The integration sites were mapped on all the human chromosomes, and a large fraction of them were identified in only one single consensus sequence read (Fig. 3C). This is probably because very few PHHs expanded following HBV DNA integration under the conditions tested, and the patterns of HBV integration remained heterogeneous.

**Figure 3.**
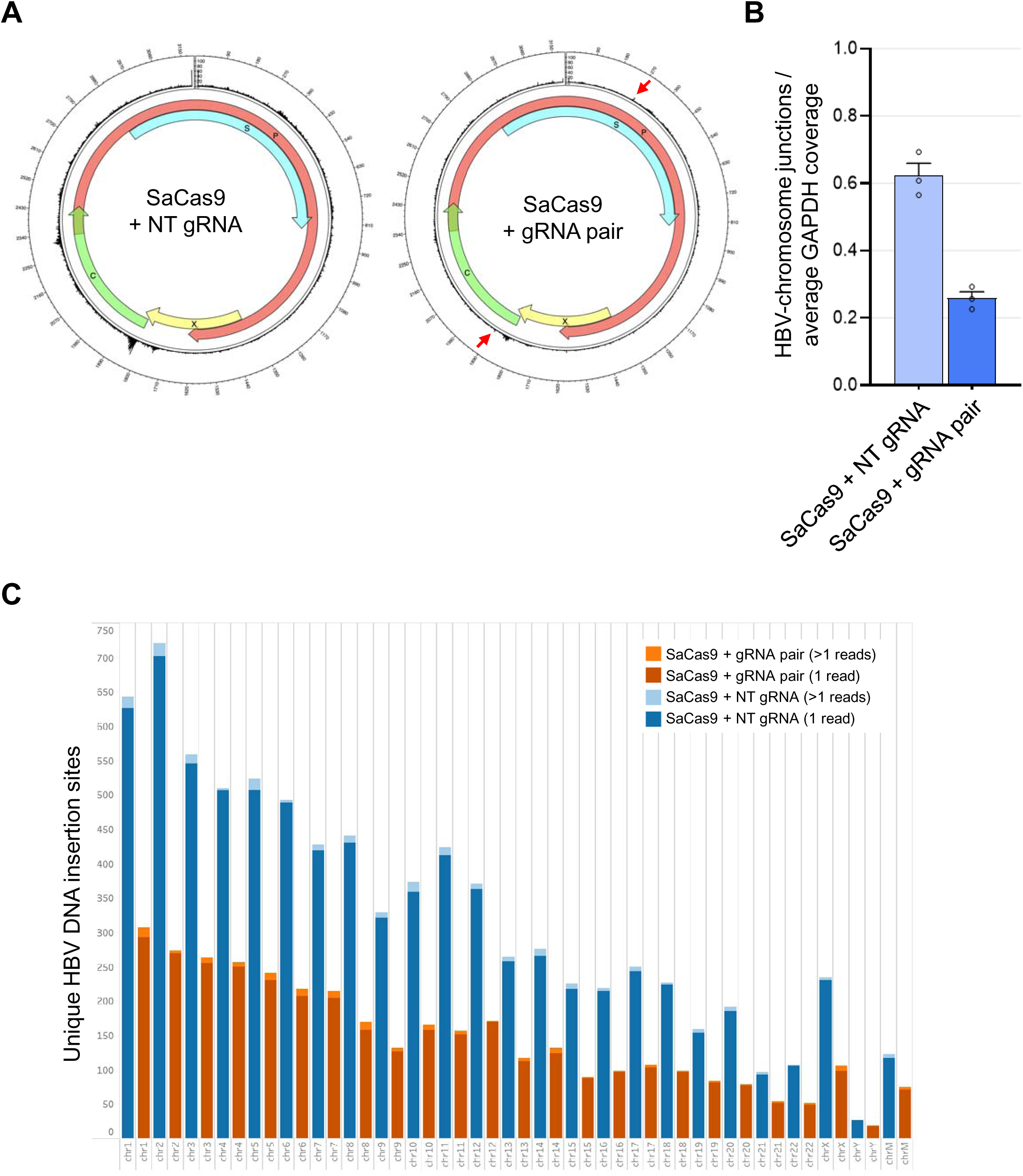
Analysis of HBV DNA insertion sites in HBV-infected PHHs by short-read hybridization capture sequencing. (A) Positions of HBV DNA connected to human chromosomes were mapped on the HBV reference genome (GenBank ID: U95551). The red arrows indicate the on-target sites. (B) The numbers of chimeric reads containing HBV- and human chromosome-derived sequences were normalized with the average sequence depth of the human GAPDH that served as an internal control. (C) The number of HBV DNA insertion sites detected in individual human chromosomes are shown. The distribution of the sites was similar between the control (blue and light blue) and edited samples (dark red and orange). The “chrM” indicates the mitochondrial DNA.

### In-vivo editing of HBV DNA in the AAV-HBV mouse model

We next studied anti-HBV activity of our editing approach in AAV-HBV mice, which were generated by transduction with a recombinant AAV8 vector carrying a 1.3-fold excess length of the genotype-D HBV genome ^31,32^. To transiently introduce anti-HBV editing in mouse hepatocytes, we intravenously administered lipid nanoparticles encapsulating SaCas9-encoding mRNA and a pair of HBV-targeting guide RNAs. We observed an LNP dose-dependent reduction in serum HBsAg, HBeAg and HBV DNA (Figs. 4A-C). To verify the molecular mechanisms underlying the viral suppression, we quantified intrahepatic HBV DNA. The assays showed that total HBV DNA significantly decreased in the livers of mice receiving the LNPs (Fig. 4D), suggesting that LNP-mediated delivery of our HBV-targeting SaCas9 inhibited HBV replication by removal of viral DNA from hepatocytes.

**Figure 4.**
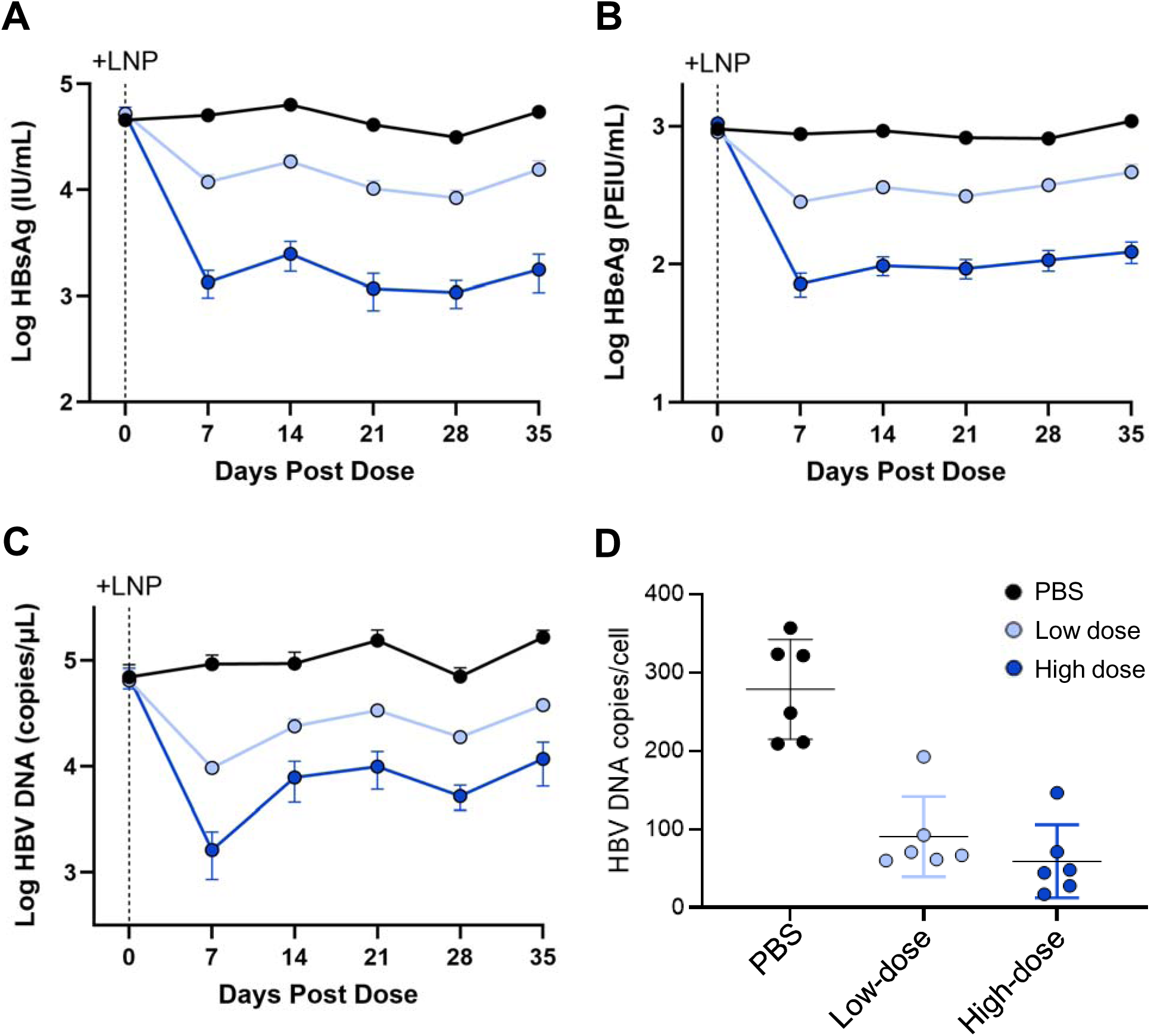
LNP-mediated delivery of the anti-HBV gene editing therapy in the AAV-HBV mouse model. (A-C) Serum levels of HBsAg (A), HBeAg (B) and HBV DNA (C) were assayed weekly following intravenous administration of PBS (control) or LNPs encapsulating the SaCas9-encoding mRNA and a pair of the HBV-targeting gRNAs at two dose levels (low dose,1.4 mg/kg; high dose, 2.9 mg/kg). (D) Total HBV DNA copies were quantified by digital PCR in liver samples harvested from each mouse 5 weeks after LNP administration.

To analyze sequence modifications of the viral DNA derived from the synthetic AAV construct and its chromosomal insertion in vivo, we sequenced by hybridization capture sequencing the viral DNA extracted from three representative liver samples selected from each of the PBS control and high-dose groups based on the copy numbers of intrahepatic HBV DNA (Supplementary Fig. 4A). The whole HBV genome was covered at 310,000 x and 210,000 x on average in the control and edited samples, respectively, and the sequence depth was greater in the redundant regions on the AAV-HBV vector (nucleotide positions 1070-1990) (Supplementary Fig. 4B). The patterns of the read coverage were slightly different between the viral DNA collected from the HBV-infected PHHs and AAV-HBV mice, suggesting that several forms of viral DNA may be present at different ratios in these two study models. Indels were detected at about 20 % of each intended target site (Fig. 5A). Excisions and inversions of sequences between the two intended target sites were also observed in 20 % and 10 % of total sequence reads spanning at least one of the two intended cut sites (Figs. 5B,C).

**Figure 5.**
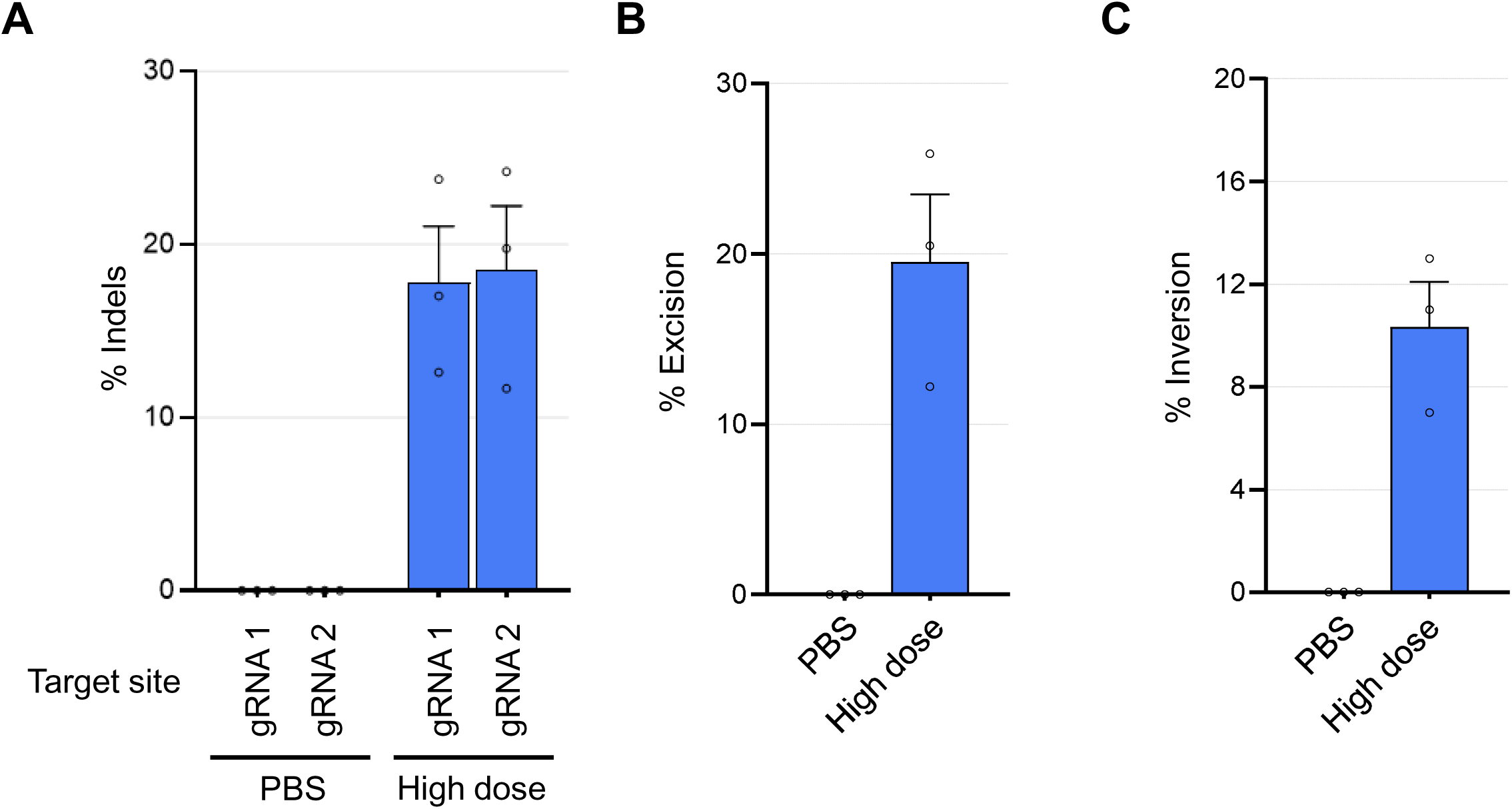
Analysis of sequence changes in HBV DNA in the AAV-HBV mouse model by short-read hybridization capture sequencing. (A) Frequency of indels at the intended target sites on HBV DNA extracted from the control and edited liver samples of the AAV-HBV. (B,C) Frequency of excisions (B) and inversions (C) of the intervening HBV sequence between the two target sites for SaCas9 gRNA 1 and SaCas9 gRNA 2. The percentages of indels and structural variants were calculated as described in the legend of Fig. 2.

The break points of HBV DNA joined to chromosomal sequences were mapped throughout its genome (Fig. 6A). Two peaks observed around DR1 in the untreated samples were proximal to the termini of dslDNA, which seems to be efficiently inserted in chromosomes of the AAV-HBV mouse model ^6,8^. Two additional spikes were found in the vicinity of the start and end positions of the 1.3 x oversized HBV genome (the positions 1070 and 1990) inserted between the AAV inverted terminal repeats (ITRs). These insertion events may be facilitated through the molecular machinery of recombinant AAV vector random integration^33^. As found in the PHHs edited with the HBV-targeting SaCas9, chimeric sequence reads containing the flanking regions of the HBV DNA on-target sites were slightly increased compared to the surrounding areas of the HBV genome in samples treated with the LNPs (Fig. 6A). The relative number of the total HBV insertion sites was reduced in edited samples when compared to the controls (Fig. 6B). Viral DNA integration sites were detected across mouse chromosomes (Fig. 6C), and many of them were supported by single consensus reads, suggesting that most hepatocytes with chromosomally integrated HBV DNA didn’t expand in the period of this experiment.

**Figure 6.**
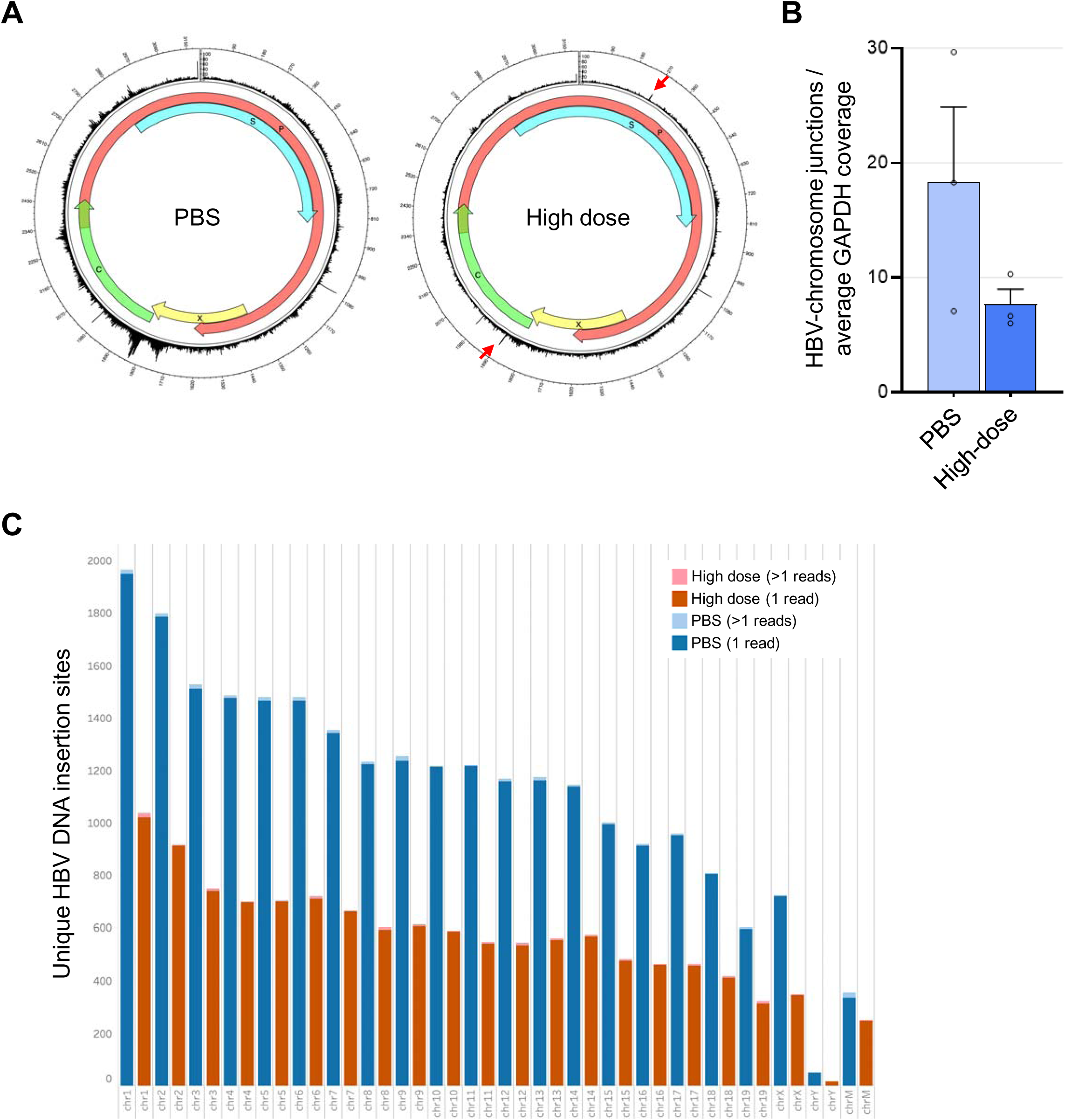
Analysis of HBV DNA insertion sites in chromosomal DNA extracted from liver samples of the AAV-HBV mouse models. (A) HBV DNA insertion sites in mouse chromosomes sequences are shown on the HBV reference genome. The red arrows indicate the on-target sites. (B) The number of sequence reads containing HBV-chromosome junctions were normalized with the average sequence coverage of the mouse Gapdh to estimate relative frequency of chromosomal HBV DNA insertion. (C) The number of HBV DNA insertion sites detected in the individual mouse chromosomes are shown. The distribution of the sites was similar between the control (blue and light blue) and edited samples (dark red and pink). The “chrM” indicates the mitochondrial DNA.

### Nuclease-mediated editing of intDNA in the transgenic HBV (Tg-HBV) mouse model

To analyze the DNA repair outcomes of double strand breaks on intDNA in vivo, we intravenously administered our anti-HBV gene editing payload encapsulated by LNPs in a Tg-HBV mouse model. This model was generated by microinjection of a synthetic DNA sequence carrying a 1.3-fold overlength of the genotype-A HBV genome, and viral transcripts from this transgenic construct were predicted to serve as the primary source of viral protein expression and replication. HBV DNA and HBsAg were secreted in blood at high levels in the hemizygous mice in a similar fashion to another transgenic animal described previously ^34,35^. Dose-dependent reduction in serum levels of HBsAg and HBV DNA and intrahepatic HBV DNA copies was observed in mice dosed with the LNPs (Figs. 7A-C). These results suggest that LNP-mediated delivery of the HBV-targeting SaCas9 efficiently targeted the chromosomally integrated HBV transgene construct and episomal viral DNAs in the Tg-HBV mice.

**Figure 7.**
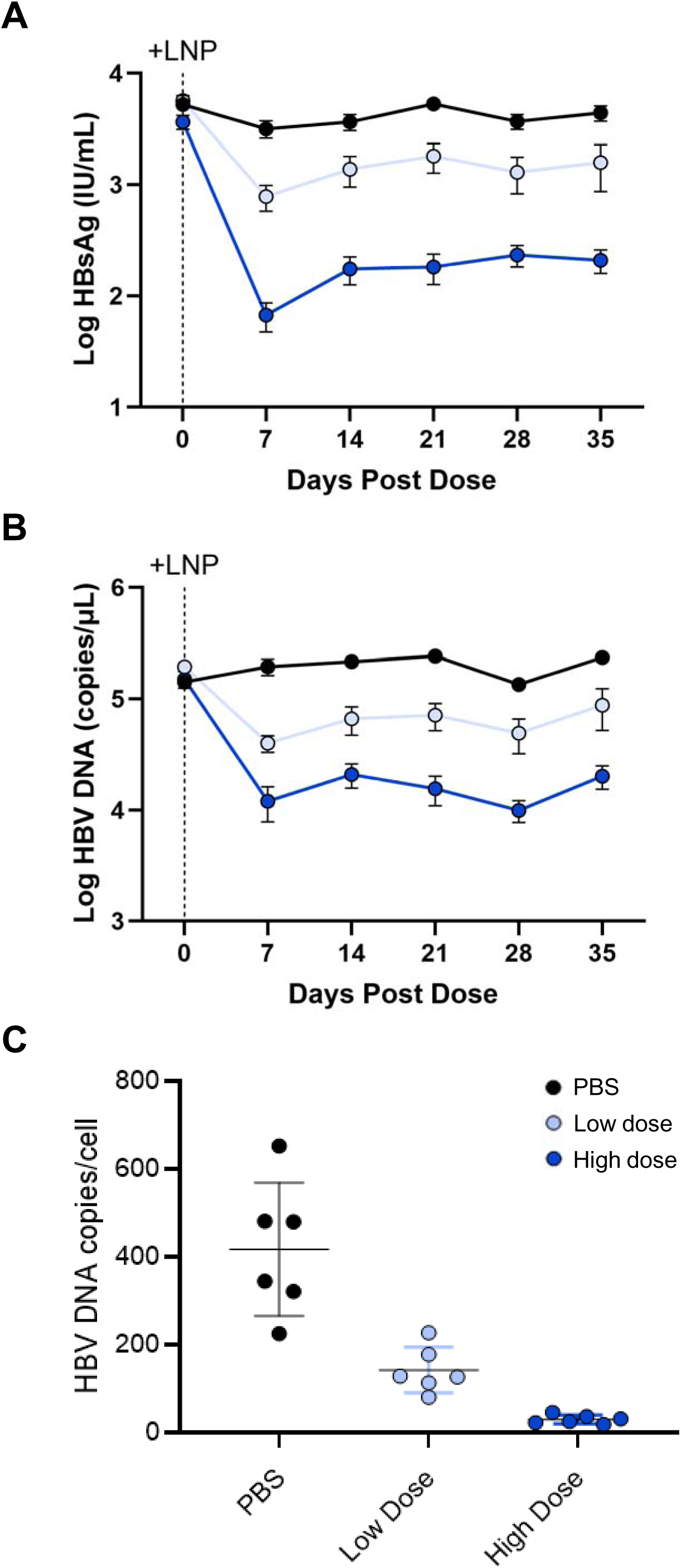
Editing by systemic LNP delivery in the Tg-HBV mouse model. (A,B) Dose-dependent reduction in serum HBsAg (A) and HBV DNA (B) was observed in the transgenic mice treated with two different quantities of LNPs encapsulating the SaCas9-encoding mRNA and a pair of the HBV-targeting gRNAs (low dose,1.6 mg/kg; high dose, 3.4 mg/kg). (C) Total HBV DNA copies were quantified by digital PCR in liver samples harvested from each mouse 5 weeks after LNP administration.

Given that all the HBV promoters were inactive in the stomach of the Tg-HBV mouse model, digital PCR should only amplify DNA from the integrated Tg-HBV construct and not from replicating HBV. We performed digital PCR on DNA extracted from the stomach tissue of one PBS treated (control) mouse, the results of which suggested that 2 copies of the full-length viral genome were inserted in each cell (Supplementary Fig. 5A). To determine the configuration of the Tg-HBV construct and its chromosomal insertion site, we performed long-read hybridization capture sequencing with DNA extracted from stomach tissues of three individual mice. The hybridization capture sequencing indicated that the transgenic construct was inserted in the reverse orientation of the mouse chromosome 3 (Supplementary Fig. 5B). It was composed of 2.3 x direct repeats of the genotype-A HBV genome containing a total of 5 target sites for the paired HBV-targeting gRNAs in addition to the partial plasmid vector backbone used for DNA cloning and 240 base pairs of HBV-derived sequence, which was in the reverse orientation to the other 2.3 x HBV genomic sequence [HBV inverted repeat (HBV-IR)]. Neither of the target sites for the paired gRNAs were present in HBV-IR.

To examine structural variants introduced by the HBV-targeting SaCas9, we analyzed HBV DNA extracted from liver samples of 5 mice per group by hybridization capture sequencing. Excisions and inversions of the intervening sequences between the 5 on-target sites were detected in the treated mice (Fig. 8A). HBV pgRNA that was transcribed from the 2.3 x oversized HBV DNA yielded rcDNA and dslDNA through reverse transcription within the viral capsid, and these viral DNA molecules can be randomly integrated into chromosomal DNA in the liver of this mouse model. As expected, insertion of HBV DNA was found at distinct chromosomal locations from that of the Tg-HBV construct. Chimeric reads representing de novo HBV DNA insertion sites were reduced by up to 80 % in the LNP-treated groups, as was previously observed in the HBV-infected PHHs and AAV-HBV mouse model (Fig. 8B). HBV-associated chromosomal translocations have been observed in liver biopsies from CHB patients ^7,36^. We identified a small number of sequence reads containing two distinct chromosomal regions interrupted by HBV DNA in both the control and treated samples. The relative frequency of chromosomal translocations involving randomly integrated HBV DNA was not substantially altered by anti-HBV multiplex gene editing (Fig. 8C). While the Tg-HBV mouse model was specifically chosen to examine editing of intDNA, we ultimately decided to exclude sequence reads containing the original Tg-HBV construct from this analysis. The structure of the Tg-HBV construct was found to be substantially different from that of integrated HBV DNA copies isolated from HBV-infected hepatocytes ^7,36^, and it is therefore difficult to conclude primary factors that contributed to chromosomal rearrangements with this synthetic DNA sequence. Taken together, these results suggest that transiently induced double strand breaks on intrahepatic HBV DNA don’t increase the risk of chromosomal aberrations caused by translocations.

**Figure 8.**
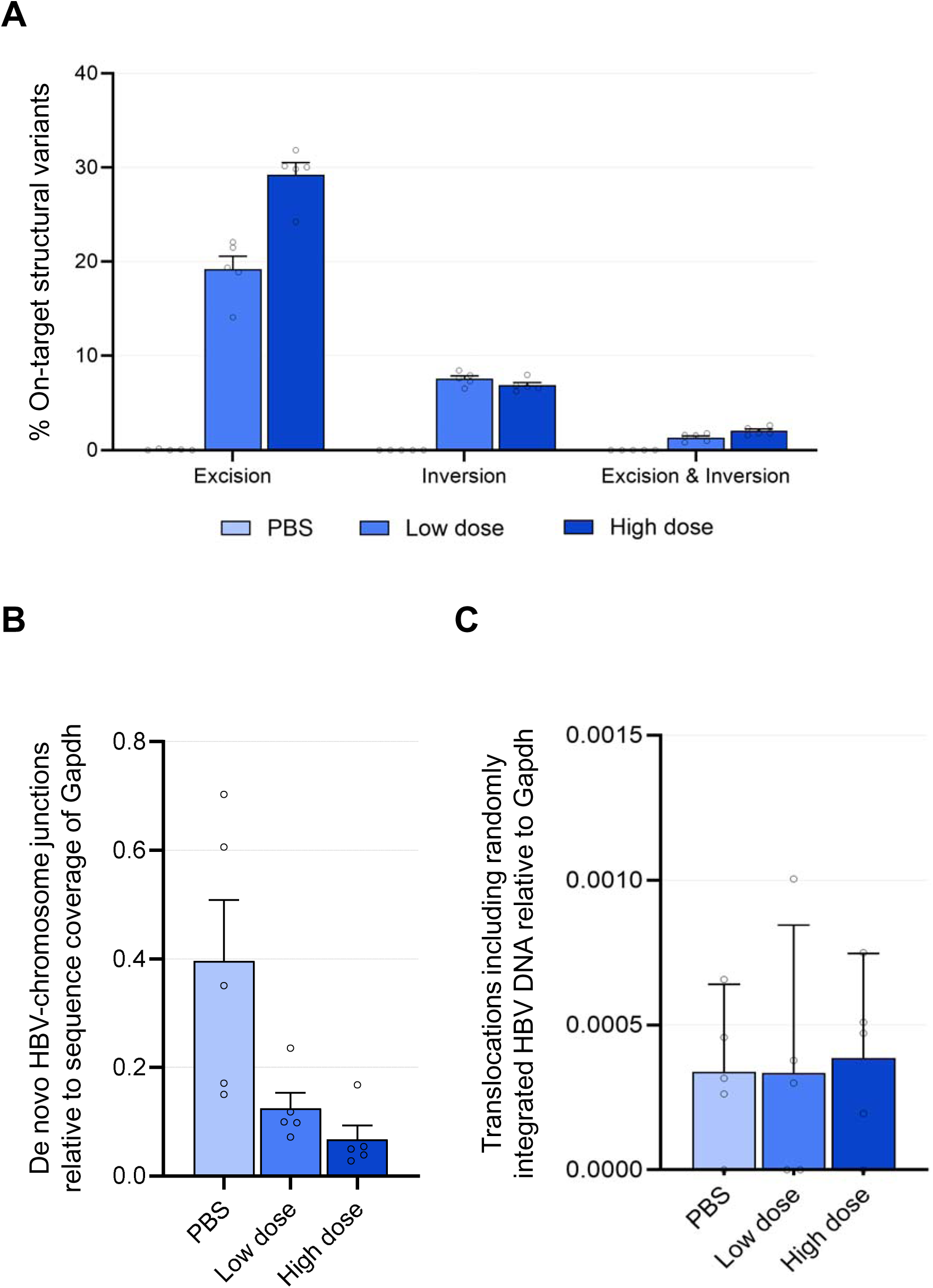
Analysis of structural variants, random HBV DNA insertions and chromosomal translocations by long-read hybridization capture sequencing. (A) The Tg-HBV mice were found to carry the 2.3 x tandem repeats of the genotype-A HBV genome in its chromosome 3. Excisions and Inversions of the chromosomally integrated HBV genomic sequences between the 5 on-target sites were quantified in liver samples from mice of the control and LNP-treated groups. Chimeric sequence reads containing de novo HBV DNA insertion sites that were distinct from the Tg-HBV construct (B) and chromosomal translocations associated with these randomly integrated HBV sequences (C) were counted and normalized with the average sequence coverage of the internal control gene locus, mouse Gapdh.

## Discussion

Treatment with NAs inhibits reverse transcription during encapsidation and significantly reduces the viral load. However, viral replication is still detected at low levels in patients who take this medication for years, leading to long-term persistence of cccDNA and slow decline of HBsAg in serum ^37^. Gene editing is a promising approach to eliminate all forms of HBV DNA including cccDNA. While cccDNA is assembled into a minichromosome along with histone and non-histone proteins ^38–40^, this nucleosome-like structure is unlikely to block access of CRISPR-Cas9 proteins to cccDNA and other forms of HBV DNA, as described in previous studies ^19–23^. In this study, we demonstrated that co-delivery of SaCas9-encoding mRNA and a pair of gRNAs we selected for targeting HBV reduced serum levels of HBsAg and HBV DNA and intrahepatic HBV DNA copies in vitro and in vivo. Either none or very low-levels of cccDNA have been detected in the AAV-HBV or Tg-HBV mouse models^31,32,34^, and viral transcripts and gene products are primarily supplied by the episomal AAV construct or chromosomally integrated HBV transgene, respectively. Recombinant AAV vectors have been thought to exist primarily as episomal forms of monomeric and concatemeric circular DNAs in a chromatin-like structure ^41^. The AAV-HBV vector that is stably maintained in mouse hepatocytes may be chromatinized in a similar fashion to HBV cccDNA. Our anti-HBV therapeutic approach, which transiently cleaves the viral target sequences, resulted in stable suppression of serum HBV biomarkers under the conditions tested. If our anti-HBV multiple gene editing fails to remove cccDNA or its seemingly equivalent AAV-HBV vector present in the animal model, viral rebound is expected to occur in a short period of time, as seen in animals that stopped receiving entecavir ^22^. Our results suggest that our HBV-targeting SaCas9 inactivates the viral DNA regardless of its forms. A liver-humanized, immunocompromised mouse is an excellent model to test treatment for CHB, where the entire steps of HBV infection are recapitulated. We previously sought to deliver mRNA to human hepatocytes of a chimeric mouse liver using conventional LNPs that adsorb serum apolipoprotein E to bind the low-density lipoprotein receptor for entry via endocytosis. We found that cellular uptake of the LNPs by human hepatocytes was inefficient in these chimeric liver animal models ^42^. We therefore chose the two other mouse models to evaluate our anti-HBV multiplex gene editing approach in vivo, although iterative HBV infection cannot be recapitulated in these animals.

To understand the outcomes of our HBV-targeting multiplex gene editing at a molecular level, we sequenced the viral and genomic DNAs from HBV-infected PHHs and the two HBV mouse models following target enrichment with hybridization capture. Detection of indels and structural variants at the HBV on-target sites indicated that SaCas9 generated double strand breaks on both episomal and integrated forms of the viral DNA. Indels and structural variants (excisions and inversions) were detected at higher frequency in the AAV-HBV mouse model than in HBV-infected PHHs. A structural discrepancy in intrahepatic HBV DNA copies is predicted between these two models and may be one of factors that impact overall DNA editing activity. A dose-dependent increase in frequency of intended structural variants was observed in the Tg-HBV mouse model, suggesting that the HBV-targeting SaCas9 could access the chromosomally integrated HBV sequence in vivo. However, we found that this transgenic construct differed significantly from typical intDNA in both size and configuration and the effects of nuclease-induced strand breaks on the integrated Tg-HBV sequence may need to be interpreted cautiously.

A previous study has shown that linearization of HBV DNA by ARCUS nucleases (which is an engineered variant of the homing endonuclease I-CreI) promotes its insertion in chromosomal sites including potential off-target sites in HBV-infected PHHs ^43^. The consequences of cutting HBV DNA seem to highly depend on the characteristics of the editing nucleases. The same research group has recently shown that the version of HBV-targeting ARCUS nucleases evaluated in preclinical development of PBGENE-HBV neither introduces off-target editing nor promotes HBV DNA integration significantly over the background ^44^. Our analysis by a series of hybridization capture sequencing experiments indicated that our anti-HBV treatment suppressed random HBV DNA integration in HBV-infected PHHs and CHB mouse models. We found that the number of break points representing insertion of dslDNA was particularly reduced as well.

Chromosomal translocations mediated by intDNA copies is another safety concern. HBV-associated chromosomal translocations have been found even in non-HCC liver biopsies from about one third of patients with CHB examined in published literature ^36^. Long-read hybridization capture sequencing suggested that chromosomal translocations involving randomly integrated HBV DNA occurred at comparable frequency in both the control and treated liver samples from the Tg-HBV mouse model. A transiently increased number of strand breaks by our therapeutic approach are unlikely to perturb the host DNA repair pathways that efficiently join DNA ends to prevent chromosomal abnormalities. However, it should be noted that no or only a few consensus sequence reads were assigned to this type of chromosomal translocation in each library replicate, and the assay procedures may need to be further improved to track extremely rare genetic changes.

Our study demonstrated that multiplex editing designed to cut intrahepatic HBV DNA substantially blocked viral transcription, protein expression and replication from the baseline. We then assessed potential safety concerns related to cleavage of episomal and integrated HBV DNA by hybridization capture sequencing. These assays indicated that our anti-HBV SaCas9 didn’t increase the risk of off-target effects in preclinical settings. Gene editing is currently the only approach to directly eliminate viral DNA that is present in patients with CHB. In addition to two classes of FDA-approved treatments, NAs and interferon α, a number of new drugs that target different aspects of the viral life cycle and host immune responses are in development ^45^. While the ideal goal of our approach is to achieve the functional cure of CHB with a monotherapy by one-time administration, repeated administration and combination therapy with other therapeutic options are also feasible and may significantly improve its clinical outcome.

## Materials and Methods

### Cell culture

The cell line 293FT was purchased from Thermo Fisher Scientific (Waltham, MA, USA), and grown in DMEM supplemented with GlutaMAX, sodium pyruvate and 10% fetal bovine serum. Primary human hepatocytes (PHHs) were purchased from BioIVT (Westbury, NY) and thawed using Cryopreserved Hepatocyte Recovery Medium and plated in William’s E Medium with Primary Hepatocyte Thawing and Plating Supplements. PHHs were cultured in William’s E Medium (Gibco) with Primary Hepatocyte Maintenance Supplements (Gibco) and 10% FBS (Gibco).

### Preparation of genomic DNA

Genomic DNA was extracted from PHH cells using a Maxwell® RSC Blood DNA Kit (Promega) with the following protocol: PHH cells were harvested, washed with PBS, then incubated at room temperature for 10 mins in 300 μL PBS with 10 μL RNase A. Then 300 μL Lysis buffer with 30 μL Proteinase K was added and samples were incubated at room temperature for 10 mins. Samples were then incubated at 80°C for 30 mins prior to loading into the Maxwell® RSC Blood DNA cartridge and ran on the Maxwell® RSC Instrument. Genomic DNA was extracted from HEK293 cells using a Maxwell® RSC Cultured Cells DNA Kit (Promega) according to the manufacturer’s protocol. Genomic DNA was extracted from mouse tissues using a MasterPure Complete DNA and RNA Purification Kit (Biosearch Technologies) with the following protocol: ∼5mg tissue samples were homogenized with a pestle in 300 µL of TE (10 mM Tris-HCl pH 8.0, 10 mM EDTA) in 1.5 mL eppendorf tubes, then 300 µL 2x Tissue and Cell Lysis Solution and 2 µL of Proteinase K was added and samples were incubated at 56°C for 1 hour. Then 2 µL RNaseA was added, and samples were incubated at 37°C for 30 mins. All other steps were performed according to the manufacturers’ protocol.

### Production of the SaCas9-encoding mRNA

The open reading frame of the SaCas9-encoding sequence optimized for human codon usage with a reduced number of thymidine bases was synthesized and inserted between the human β-globin 5’ UTR and 3’ UTR connected to a polyadenine tail flanked by BspQI target sites of a standard cloning vector. In-vitro transcription by the T7 RNA polymerase was performed using the BspQI-linearized plasmid and N1-methylpseudouridine-5’-triphosphate instead of uridine-5’-triphosphate, and the cap-1 structure was added with the vaccinia capping enzyme and 2’-O-methyltransferase.

### Generation of the HBV reporter cell line

The HBV reporter cell line was generated as described previously ^15^. Briefly, the synthetic sequence illustrated in Supplementary Fig. 1B was synthesized and cloned in the pUC57-Kan vector at GenScript. The enhanced green fluorescent protein (EGFP) gene fused to the puromycin-resistant gene via the self-cleaving peptide P2A-coding sequence was under the control of the murine stem cell virus promoter with a reduced number of CpG sites ^46^, and transcription of the downstream sequence including the synthetic exons and the blue fluorescent protein (BFP) and blasticidin-S deaminase genes was activated by the human EF1α core promoter. The entire sequence was flanked by the two AAVS1 CRISPR-Cas9 target to be linearized for targeted knock-in with Streptococcus pyogenes Cas9 ribonucleoprotein complex (SpCas9 RNP). Five hundred thousand 293FT cells were transfected with 0.5 μg of the reporter construct and 20 pmol of SpCas9 RNP by electroporation using 4D-Nucleofector System (Lonza) in 20 μl of the SF Cell Line Nucleofector Solution under the CM-130 program. Cells with the chromosomally integrated reporter construct were selected in the presence of 2 μg/ml puromycin, and single clones were isolated by limiting dilution. Biallelic knock-in was confirmed by multiplex digital PCR.

### Evaluation of intended editing in the HBV reporter cell line

To quantify excision and indels, 1.2 × 10^5^ HBV reporter cells were seeded per well in a 24-well plate and transfected with 0.5 μg each of the SaCas9-encoding mRNA and a pair of HBV-targeting guide RNAs using Lipofectamine MessengerMAX. The cells were harvested 5 days post transfection and genomic DNA was extracted. To calculate frequency of excision, the intervening sequence between the two on-target sites and the downstream sequence encoding mTagBFP2 were quantified by multiplex digital PCR. Indels at the target sites that were not removed by excision were assayed by Sanger sequencing of PCR-amplified fragments.

### GUIDE-seq

GUIDE-seq was conducted, and sequencing data was analyzed following the procedures described previously with minor modifications ^15,26^. Briefly, the HBV reporter cell line (200,000 cells) was transfected with 1 μg each of the SaCas9-encoding mRNA and one of the two gRNAs and 15 pmol of the double stranded oligonucleotide (dsODN) by electroporation using 4D-Nucleofector System (Lonza) in 20 μl of the SF Cell Line Nucleofector Solution under the CM-130 program. The cells were harvested 4 days post transfection for DNA extraction followed by library preparation.

### Targeted amplicon sequencing

To quantify indels at the nominated sites identified by GUIDE-seq, the HBV reporter cell line (200,000 cells) was transfected with 1 μg each of the SaCas9-encoding mRNA and either of the two gRNAs by electroporation using 4D-Nucleofector System (Lonza) in 20 μl of the SF Cell Line Nucleofector Solution under the CM-130 program. Sequences spanning the HBV on-target sies and nominated sites selected by GUIDE-seq were PCR-amplified using Q5 High-Fidelity 2X Master Mix (New England Biolabs) and sequenced using a 2 × 150-bp paired-end run on the MiSeq system (Illumina). Targeted amplicon sequencing analysis was performed using CRISPResso2 ^47^ as described previously ^15^.

### Hybridization capture sequencing

Library preparation and target capture for HBV sequences were performed using SureSelect XT HS2 reagents (Agilent Technologies) according to the manufacturer’s protocol. Capture probes were designed to target 2 HBV strains (GenBank accession numbers: U95551 and AY128092) and GAPDH. The captured libraries were pooled and sequenced by Novogene using a 2 × 150-bp paired-end run on the NovaSeq X Plus platform (Illumina). Long-read hybridization capture sequencing was performed according to Agilent’s Target Capture Long-Read Sequencing Using the Agilent SureSelect XT HS2 Target Enrichment System application note with the following modifications. After captured-library amplification, 49 μL of each library was used as input for steps 4 through 7 of PacBio’s Preparing whole genome libraries using the HiFi prep kit 96 protocol. The nuclease-treated libraries were then cleaned up an additional time using the SMRTbell cleanup beads and eluted in 26 μL. The purified libraries were then sent to NovoGene for step 9 (Annealing, binding, and cleanup) of the PacBio protocol, followed by library pooling and sequencing on the PacBio Revio system. The capture probes used were the same as described above.

### HBV infection and transfection of PHHs

Stock vials of HBV genotype D produced from HepG2-derived cell line, HepAD38 was purchased from ImQuest BioSciences. Primary human hepatocytes obtained from BioIVT were seeded at 3.5 × 10^5^ live cells per well in a 24-well plate. After 24 hours, cells were inoculated with HBV at 500 genome equivalents per cell in the presence of 4% polyethylene glycol 8000 (PEG 8000) (Sigma-Aldrich) and 2% DMSO (Corning) at 37°C for 16 hours. After 4 days, cells were transfected with 0.5 μg of the SaCas9-encoding mRNA and 0.25 µg each of a pair of HBV-targeting guide RNAs using Lipofectamine MessengerMAX. After 4 days and 7 days, supernatants were collected, and cells were harvested for gDNA extraction.

### HBsAg and HBeAg analysis

HBsAg and HBeAg levels in cell culture medium were measured with HBeAg- and HBsAg-specific chemiluminescence assays (CLIA) (Ig Biotechnology). Fifty microliters (50 µL) of culture supernatant and standard curve calibrators were incubated with 50 µL enzyme conjugate (HRP labeled anti-HBs or anti-HBe MAb in PBS containing Casein, BSA, and ProlClin 300) in a 96-well test plate, shaken on a plate shaker for 60 seconds and then incubated for 60 minutes at 37°C. Following incubation, the contents of the plate were removed, and the plate was washed six times with 350 µL of Wash Solution (PBS-Tween, 1X). Substrate solution (hydrogen peroxide in buffer solution) was prepared and 50 µL was added to each well. The plate was incubated for 10 minutes in the dark, then read on a FlexStation 3 plate reader (Molecular Devices) measuring the Relative Light Units (RLUs) of each well. From the RLUs, the amount of HBeAg (PEIU/mL) or HBsAg (IU/mL) in each sample was determined.

### LNP formulation

The SaCas9-encoding mRNA and a pair of the gRNAs were mixed at 2:1:1 (in weight) and formulated in four lipid components using a microfluidic instrument at the N/P ratio of 6:1. For the AAV-HBV mouse model, SM-102, DSPC, cholesterol and DMG-PEG2000 dissolved in ethanol at 50:10:38.5:1.5 (in molar) were used. The aqueous phase was substituted with 1 x PBS using a centrifugal filter. After concentration, the LNPs were sterilized through a 0.2-µm filter. For the Tg-HBV mouse model, the lipid solution containing LP-01, DSPC, cholesterol and DMG-PEG2000 at 50:9:38.5:3 (in molar) was used to prepare LNPs. After dialysis in 50 mM Tris and 45 mM NaCl, pH 7.5 overnight, the LNPs were concentrated using a centrifugal filter and sterilized through a 0.2-µm filter. Sucrose was added to 5 % (v/v) before storage at −80 °C.

### Ethic statement and animal care

All mice were housed in individually ventilated cages at the WuXi AppTec’s animal facility in Shanghai, China. Animal care and euthanasia for the studies with the AAV-HBV and Tg-HBV mouse models were conducted following standard operating procedures approved by the WuXi Institutional Animal Care and Use Committee (IACUC; protocol numbers ID01-013-2021v1.0 and ID01-SH002-2023v1.2). The animal facility and IACUC were fully accredited by Association of Assessment and Accreditation of Laboratory Animal Care International (AAALAC).

### In-vivo editing in the AAV-HBV mouse model

Each of the 5-6 weeks old C57BL/6J mice were transduced with 1 × 10^10^ vg of the AAV-HBV vector by tail vein injection. Serum samples were collected by submandibular bleeding to assay serum level of HBV DNA, HBsAg and HBeAg weekly from 3 weeks post the AAV administration until termination of the in-life stage. HBV DNA was isolated with QIAamp 96 Blood DNA Kit (Qiagen) following the manufacturer’s instructions and its copy number was determined by quantitative PCR using TaqMan Universal PCR Master Mix (ThermoFisher Scientific). HBsAg and HBeAg were assayed using HBsAg and HBeAg ELISA kits (Autobio). Mice were divided into 3 groups to minimize differences in body weights and serum HBV DNA, HBsAg and HBeAg among the groups. PBS or 35 µg or 70 µg of the anti-HBV gene editing payload encapsulated by the LNPs were administered into mice of each group. All the mice were euthanized 5 weeks after LNP administration, and the partial left lobe of liver from each mouse was harvested, flash-frozen in liquid nitrogen, and stored at −80 °C until use.

### In-vivo editing in the Tg-HBV mouse model

Seven-week-old Tg-HBV female mice were purchased from Beiing Vitalstar Biotechnology Co.. Ltd. Each of 5-6 weeks old. This mouse model was generated through pronuclear microinjection of the linearized DNA fragment carrying the 1.3-fold oversized HBV genome (GenBank ID: AF305422.1, subtype adw2, genotype A) into the fertilized egg of a C57BL/6 mouse. Serum samples were collected by submandibular bleeding to assay serum level of HBV DNA, HBsAg and HBeAg one week prior to LNP administration. HBV DNA, HBsAg and HBeAg were quantified as described above. The assay results were used to divide the mice into 3 groups for equalizing body weights and serum HBV DNA and HBsAg among the groups. PBS or 30 µg or 60 µg of the anti-HBV gene editing payload encapsulated by the LNPs were administered into mice of each group by tail vein injection at 9 weeks old. Each mouse was weekly bled to assay serum HBV DNA, HBsAg and HBeAg. All the mice were euthanized 5 weeks after LNP administration, and tissue samples including the partial left lobe of liver and part of stomach from each mouse were harvested, flash-frozen in liquid nitrogen, and stored at −80 °C until use.

### In-silico HBV viral target site search

HBV viral sequences were downloaded from HBVdb ^25^ for genotypes A to H. Customized python script was used to search for target site with length of 21 nt and PAM sequence of NNGRRT. Target sites from all HBV sequences were compiled, and the occurrence of each target site was computed within each genotype (Sup. Fig 1A). The occurrence of a target site represents the percentage of genomes within a certain genotype containing that target site. Those with over 80% occurrences across all genotypes were selected for downstream analysis.

### Homologous site search in the human reference genome

Human genomic sites that shared high sequence identity with the selected gRNA on-target sequences were identified using an Nextflow pipeline as described previously ^15^. Briefly, CasOffinder ^48^ was used to search for homologous sites in human genome (hg38_analysis_set) with PAM of NNNNNN and length of 21 nt allowing up to 6 total differences and 2 gaps. Duplicated sites were identified by sharing the same predicted PAM locations and removed from the result. For each unique predicted PAM location, the homologous site in human genome with minimal number of mismatches and gaps was kept. Mismatches within PAM region against NNGRRT were calculated and added to the total number of mismatches. The sites with up to 6 total differences are shown in Supplementary Table 1.

### Data analysis for short-read hybridization capture sequencing with DNA samples with HBV-infected PHHs and AAV-HBV mice

Sequence reads obtained from short-read hybridization capture analysis were analyzed using a customized Nextflow pipeline developed in house. Demultiplexed sequencing reads were first preprocessed and trimmed for sequencing adapters using FASTP ^49^. UMI sequences were extracted, and reads were collapsed into UMI consensus sequences using FGBIO recommended pipeline ^50^. Editing frequencies around gRNA target sites were analyzed and quantified using CRISPResso2 ^47^. Structural variants were identified using FGSV ^51^. Reads with excision or inversion were identified if DNA break point was within 25 nt of both gRNA cut sites. Sequencing depth at regions of interest were computed using Mosdepth ^52^. Host genomic sites with potential HBV integration were searched for homology against each gRNA sequence using Calitas ^53^ with ±60 bp window. Reference genome used during this analysis include human genome (hg38 analysis set), mouse genome (mm39), HBV reference AYW (GenBank: U95551.1), and AAV2 ITR sequence.

### Data analysis for long-read hybridization capture sequencing with DNA samples from Tg-HBV mice

Sequence reads obtained by long-read hybridization capture sequencing were analyzed using a customized Nextflow pipeline developed in house. Briefly, demultiplexed PacBio sequencing reads were quality checked using NanoPlot ^54^ and pre-processed to remove Illumina sequencing adapters using LIMA ^55^. Duplicated sequencing reads were identified and removed using Pbmarkdup ^56^. 3bp from each end of sequencing reads were extracted as UMI sequences. Reads were aligned to reference genomes using bwa ^57^ and ngm_lr ^58^. Reference genomes include mouse genome (mm39) and HBV reference AYW (GenBank: U95551.1), and pK18msr (GenBank: AB694752.1). Sequencing depth of region of interested was computed using bwa mapped reads with Mosdepth ^52^. Reads aligned to HBV reference genome were extracted and further characterized for read composition. HBV containing reads were blasted against the same reference genome set using command line blast+ ^59^. Customized python script was used to remove duplicated reads sharing the same UMI sequences. Reads with potential integration or translocation were identified by having sequencing from host genome wrapping around HBV sequences, gaps between host sequence and HBV sequence less than 50 nt, and blast identity greater than or equal to 95%. The host genomic sites with potential HBV integration or translocation were searched for sequence homology against gRNAs using Calitas ^53^ with ±60 bp window. Customized python scripts were used to quantify reads with excision or inversion with ±20 nt window near gRNA target sites.

### Analysis of results from GUIDE-seq

GUIDE-seq analysis was performed using in house pipeline as described previously ^15^. Briefly, sequencing reads were demultiplexed using deML ^60^. Identification of dsODN integration sites was modified from previous publication ^26^. Genomic sites with potential dsODN integration were searched for sequence homology against gRNA using Calitas ^53^ with ±60 bp window. Mosdepth tool was used to calculate base coverage at the identified sites. Genomic sites with total number of differences less than or equal to 7 in the alignment with the HBV on-target sites were selected for downstream analysis.

## References

1 Mendenhall, M. A., Hong, X. & Hu, J. Hepatitis B Virus Capsid: The Core in Productive Entry and Covalently Closed Circular DNA Formation. Viruses 15, doi:10.3390/v15030642 (2023).

2 Revill, P. A. et al. A global scientific strategy to cure hepatitis B. Lancet Gastroenterol Hepatol 4, 545–558, doi:10.1016/S2468-1253(19)30119-0 (2019).

3 Foundation, H. B. Acute vs. Chronic Hepatitis B, <https://www.hepb.org/what-is-hepatitis-b/what-is-hepb/acute-vs-chronic/>

4 Sung, W. K. et al. Genome-wide survey of recurrent HBV integration in hepatocellular carcinoma. Nat Genet 44, 765–769, doi:10.1038/ng.2295 (2012).

5 Zhao, L. H. et al. Genomic and oncogenic preference of HBV integration in hepatocellular carcinoma. Nat Commun 7, 12992, doi:10.1038/ncomms12992 (2016).

6 Tu, T., Budzinska, M. A., Vondran, F. W. R., Shackel, N. A. & Urban, S. Hepatitis B Virus DNA Integration Occurs Early in the Viral Life Cycle in an In Vitro Infection Model via Sodium Taurocholate Cotransporting Polypeptide-Dependent Uptake of Enveloped Virus Particles. J Virol 92, doi:10.1128/JVI.02007-17 (2018).

7 Zoulim, F., Chen, P. J., Dandri, M., Kennedy, P. T. & Seeger, C. Hepatitis B virus DNA integration: Implications for diagnostics, therapy, and outcome. J Hepatol 81, 1087–1099, doi:10.1016/j.jhep.2024.06.037 (2024).

8 Li, X. et al. The function of targeted host genes determines the oncogenicity of HBV integration in hepatocellular carcinoma. J Hepatol 60, 975–984, doi:10.1016/j.jhep.2013.12.014 (2014).

9 Yang, W. & Summers, J. Integration of hepadnavirus DNA in infected liver: evidence for a linear precursor. J Virol 73, 9710–9717, doi:10.1128/JVI.73.12.9710-9717.1999 (1999).

10 Sun, Y. et al. Persistent Low Level of Hepatitis B Virus Promotes Fibrosis Progression During Therapy. Clin Gastroenterol Hepatol 18, 2582–2591 e2586, doi:10.1016/j.cgh.2020.03.001 (2020).

11 Broquetas, T. & Carrion, J. A. Current Perspectives on Nucleos(t)ide Analogue Therapy for the Long-Term Treatment of Hepatitis B Virus. Hepat Med 14, 87–100, doi:10.2147/HMER.S291976 (2022).

12 Ghany, M. G., Buti, M., Lampertico, P., Lee, H. M. & Faculty, A.-E. H.-H. T. E. C. Guidance on treatment endpoints and study design for clinical trials aiming to achieve cure in chronic hepatitis B and D: Report from the 2022 AASLD-EASL HBV-HDV Treatment Endpoints Conference. Hepatology 78, 1654–1673, doi:10.1097/HEP.0000000000000431 (2023).

13 Ghany, M. G., Buti, M., Lampertico, P., Lee, H. M. & Faculty, A.-E. H.-H. T. E. C. Guidance on treatment endpoints and study design for clinical trials aiming to achieve cure in chronic hepatitis B and D: Report from the 2022 AASLD-EASL HBV-HDV Treatment Endpoints Conference. J Hepatol 79, 1254–1269, doi:10.1016/j.jhep.2023.06.002 (2023).

14 Dash, P. K. et al. Sequential LASER ART and CRISPR Treatments Eliminate HIV-1 in a Subset of Infected Humanized Mice. Nat Commun 10, 2753, doi:10.1038/s41467-019-10366-y (2019).

15 Amrani, N. et al. CRISPR-Cas9-mediated genome editing delivered by a single AAV9 vector inhibits HSV-1 reactivation in a latent rabbit keratitis model. Mol Ther Methods Clin Dev 32, 101303, doi:10.1016/j.omtm.2024.101303 (2024).

16 Gillmore, J. D. et al. CRISPR-Cas9 In Vivo Gene Editing for Transthyretin Amyloidosis. N Engl J Med 385, 493–502, doi:10.1056/NEJMoa2107454 (2021).

17 Musunuru, K. et al. In vivo CRISPR base editing of PCSK9 durably lowers cholesterol in primates. Nature 593, 429–434, doi:10.1038/s41586-021-03534-y (2021).

18 Li, H. et al. Inhibition of HBV Expression in HBV Transgenic Mice Using AAV-Delivered CRISPR-SaCas9. Front Immunol 9, 2080, doi:10.3389/fimmu.2018.02080 (2018).

19 Kayesh, M. E. H. et al. Development of an in vivo delivery system for CRISPR/Cas9-mediated targeting of hepatitis B virus cccDNA. Virus Res 290, 198191, doi:10.1016/j.virusres.2020.198191 (2020).

20 Stone, D. et al. CRISPR-Cas9 gene editing of hepatitis B virus in chronically infected humanized mice. Mol Ther Methods Clin Dev 20, 258–275, doi:10.1016/j.omtm.2020.11.014 (2021).

21 Martinez, M. G. et al. CRISPR-Cas9 Targeting of Hepatitis B Virus Covalently Closed Circular DNA Generates Transcriptionally Active Episomal Variants. mBio 13, e0288821, doi:10.1128/mbio.02888-21 (2022).

22 Smekalova, E. M. et al. Cytosine base editing inhibits hepatitis B virus replication and reduces HBsAg expression in vitro and in vivo. Mol Ther Nucleic Acids 35, 102112, doi:10.1016/j.omtn.2023.102112 (2024).

23 Yao, Z. Q. et al. The potential of HBV cure: an overview of CRISPR-mediated HBV gene disruption. Front Genome Ed 6, 1467449, doi:10.3389/fgeed.2024.1467449 (2024).

24 Araujo, N. M., Teles, S. A. & Spitz, N. Comprehensive Analysis of Clinically Significant Hepatitis B Virus Mutations in Relation to Genotype, Subgenotype and Geographic Region. Front Microbiol 11, 616023, doi:10.3389/fmicb.2020.616023 (2020).

25 Hayer, J. et al. HBVdb: a knowledge database for Hepatitis B Virus. Nucleic Acids Res 41, D566–570, doi:10.1093/nar/gks1022 (2013).

26 Tsai, S. Q. et al. GUIDE-seq enables genome-wide profiling of off-target cleavage by CRISPR-Cas nucleases. Nat Biotechnol 33, 187–197, doi:10.1038/nbt.3117 (2015).

27 Zhu, L. J., et al. GUIDEseq: a bioconductor package to analyze GUIDE-Seq datasets for CRISPR-Cas nucleases. BMC Genomics 18, 379, doi:10.1186/s12864-017-3746-y (2017).

28 Tycko, J. et al. Pairwise library screen systematically interrogates Staphylococcus aureus Cas9 specificity in human cells. Nat Commun 9, 2962, doi:10.1038/s41467-018-05391-2 (2018).

29 Jiang, S. et al. Re-evaluation of the carcinogenic significance of hepatitis B virus integration in hepatocarcinogenesis. PLoS One 7, e40363, doi:10.1371/journal.pone.0040363 (2012).

30 Gong, S. S., Jensen, A. D., Chang, C. J. & Rogler, C. E. Double-stranded linear duck hepatitis B virus (DHBV) stably integrates at a higher frequency than wild-type DHBV in LMH chicken hepatoma cells. J Virol 73, 1492–1502, doi:10.1128/JVI.73.2.1492-1502.1999 (1999).

31 Lucifora, J. et al. Detection of the hepatitis B virus (HBV) covalently-closed-circular DNA (cccDNA) in mice transduced with a recombinant AAV-HBV vector. Antiviral Res 145, 14–19, doi:10.1016/j.antiviral.2017.07.006 (2017).

32 Du, Y. et al. In Vivo Mouse Models for Hepatitis B Virus Infection and Their Application. Front Immunol 12, 766534, doi:10.3389/fimmu.2021.766534 (2021).

33 McCarty, D. M., Young, S. M., Jr. & Samulski, R. J. Integration of adeno-associated virus (AAV) and recombinant AAV vectors. Annu Rev Genet 38, 819–845, doi:10.1146/annurev.genet.37.110801.143717 (2004).

34 Guidotti, L. G., Matzke, B., Schaller, H. & Chisari, F. V. High-level hepatitis B virus replication in transgenic mice. J Virol 69, 6158–6169, doi:10.1128/JVI.69.10.6158-6169.1995 (1995).

35 Fumagalli, V. et al. Serum HBsAg clearance has minimal impact on CD8+ T cell responses in mouse models of HBV infection. J Exp Med 217, doi:10.1084/jem.20200298 (2020).

36 van Buuren, N. et al. Targeted long-read sequencing reveals clonally expanded HBV-associated chromosomal translocations in patients with chronic hepatitis B. JHEP Rep 4, 100449, doi:10.1016/j.jhepr.2022.100449 (2022).

37 Yu, T. et al. Evidence of Residual Ongoing Viral Replication in Chronic Hepatitis B Patients Successfully Treated With Nucleos(t)ide Analogues. J Infect Dis 227, 675–685, doi:10.1093/infdis/jiac493 (2023).

38 Bock, C. T. et al. Structural organization of the hepatitis B virus minichromosome. J Mol Biol 307, 183–196, doi:10.1006/jmbi.2000.4481 (2001).

39 Pollicino, T. et al. Hepatitis B virus replication is regulated by the acetylation status of hepatitis B virus cccDNA-bound H3 and H4 histones. Gastroenterology 130, 823–837, doi:10.1053/j.gastro.2006.01.001 (2006).

40 Tropberger, P. et al. Mapping of histone modifications in episomal HBV cccDNA uncovers an unusual chromatin organization amenable to epigenetic manipulation. Proc Natl Acad Sci U S A 112, E5715–5724, doi:10.1073/pnas.1518090112 (2015).

41 Penaud-Budloo, M. et al. Adeno-associated virus vector genomes persist as episomal chromatin in primate muscle. J Virol 82, 7875–7885, doi:10.1128/JVI.00649-08 (2008).

42 Stone, D. et al. Serum factors create species-specific barriers to hepatic gene transfer by lipid nanoparticles in liver-humanized mice. Molecular Therapy Methods & Clinical Development, doi:10.1016/j.omtm.2025.101470 (2025).

43 Gorsuch, C. L. et al. Targeting the hepatitis B cccDNA with a sequence-specific ARCUS nuclease to eliminate hepatitis B virus in vivo. Mol Ther 30, 2909–2922, doi:10.1016/j.ymthe.2022.05.013 (2022).

44 Harrison, E. et al. LBP-022 Preclinical safety data for PBGENE-HBV gene editing program supports advancement to clinical trials as a potentially curative treatment for chronic hepatitis B. Journal of Hepatology 80, doi:10.1016/S0168-8278(24)00589-0 (2024).

45 Foundation, H. B. Drug Watch, <https://www.hepb.org/treatment-and-management/drug-watch/> (2025).

46 Dickson, R. A. The scientific basis of treatment of idiopathic thoracic scoliosis. Acta Orthop Belg 58 Suppl 1, 107–110 (1992).

47 Pinello, L. et al. Analyzing CRISPR genome-editing experiments with CRISPResso. Nat Biotechnol 34, 695–697, doi:10.1038/nbt.3583 (2016).

48 Bae, S., Park, J. & Kim, J. S. Cas-OFFinder: a fast and versatile algorithm that searches for potential off-target sites of Cas9 RNA-guided endonucleases. Bioinformatics 30, 1473–1475, doi:10.1093/bioinformatics/btu048 (2014).

49 Chen, S., Zhou, Y., Chen, Y. & Gu, J. fastp: an ultra-fast all-in-one FASTQ preprocessor. Bioinformatics 34, i884–i890, doi:10.1093/bioinformatics/bty560 (2018).

50 Fennell, T. H., N. fgbio Best Practise FASTQ -> Consensus Pipeline, <https://github.com/fulcrumgenomics/fgbio/blob/main/docs/best-practice-consensus-pipeline.md>

51 Fulcrum. fgsv, <https://github.com/fulcrumgenomics/fgsv>

52 Pedersen, B. S. & Quinlan, A. R. Mosdepth: quick coverage calculation for genomes and exomes. Bioinformatics 34, 867–868, doi:10.1093/bioinformatics/btx699 (2018).

53 Fennell, T. et al. CALITAS: A CRISPR-Cas-aware ALigner for In silico off-TArget Search. CRISPR J 4, 264–274, doi:10.1089/crispr.2020.0036 (2021).

54 De Coster, W. & Rademakers, R. NanoPack2: population-scale evaluation of long-read sequencing data. Bioinformatics 39, doi:10.1093/bioinformatics/btad311 (2023).

55 PacBio. Lima, <https://lima.how/>

56 PacBio. pbmarkdup, <https://github.com/PacificBiosciences/pbmarkdup> (

57 Jung, Y. & Han, D. BWA-MEME: BWA-MEM emulated with a machine learning approach. Bioinformatics 38, 2404–2413, doi:10.1093/bioinformatics/btac137 (2022).

58 Sedlazeck, F. J. et al. Accurate detection of complex structural variations using single-molecule sequencing. Nat Methods 15, 461–468, doi:10.1038/s41592-018-0001-7 (2018).

59 Camacho, C. et al. BLAST+: architecture and applications. BMC Bioinformatics 10, 421, doi:10.1186/1471-2105-10-421 (2009).

60 Renaud, G., Stenzel, U., Maricic, T., Wiebe, V. & Kelso, J. deML: robust demultiplexing of Illumina sequences using a likelihood-based approach. Bioinformatics 31, 770–772, doi:10.1093/bioinformatics/btu719 (2015).

