## Supplementary Table 1 for "Multiplex Gene Editing Suppresses Random Integration of Hepatitis B Virus DNA in Chronically Infected Liver"

**Supplementary Table 1. Human genomic sites that are homologous to the viral target sites for SaCas9 gRNA 1 and SaCas9 gRNA 2.**

| SaCas9 gRNA 1 | | |
|---|---|---|
| # Mismatch | # Gap | # Site |
| 3 | 0 | 3 |
| 4 | 0 | 29 |
| 5 | 0 | 310 |
| 6 | 0 | 2630 |
| 1 | 1 | 1 |
| 2 | 1 | 3 |
| 3 | 1 | 85 |
| 4 | 1 | 1027 |
| 5 | 1 | 11244 |
| 2 | 2 | 3 |
| 3 | 2 | 170 |
| 4 | 2 | 1949 |

| SaCas9 gRNA 2 | | |
|---|---|---|
| # Mismatch | # Gap | # Site |
| 3 | 0 | 0 |
| 4 | 0 | 20 |
| 5 | 0 | 187 |
| 6 | 0 | 1794 |
| 1 | 1 | 0 |
| 2 | 1 | 3 |
| 3 | 1 | 48 |
| 4 | 1 | 612 |
| 5 | 1 | 6629 |
| 2 | 2 | 9 |
| 3 | 2 | 148 |
| 4 | 2 | 1861 |
