## Supplementary Figures for "Multiplex Gene Editing Suppresses Random Integration of Hepatitis B Virus DNA in Chronically Infected Liver"

**A**

| Guide Name | HBV genotype A | B | C | D | E | F | G | H |
|---|---|---|---|---|---|---|---|---|
| SaCas9 gRNA 1 | 95.0 | 87.2 | 95.7 | 94.9 | 97.2 | 96.9 | 100.0 | 100.0 |
| SaCas9 gRNA 2 | 96.2 | 97.4 | 98.4 | 98.1 | 97.2 | 99.0 | 100.0 | 100.0 |

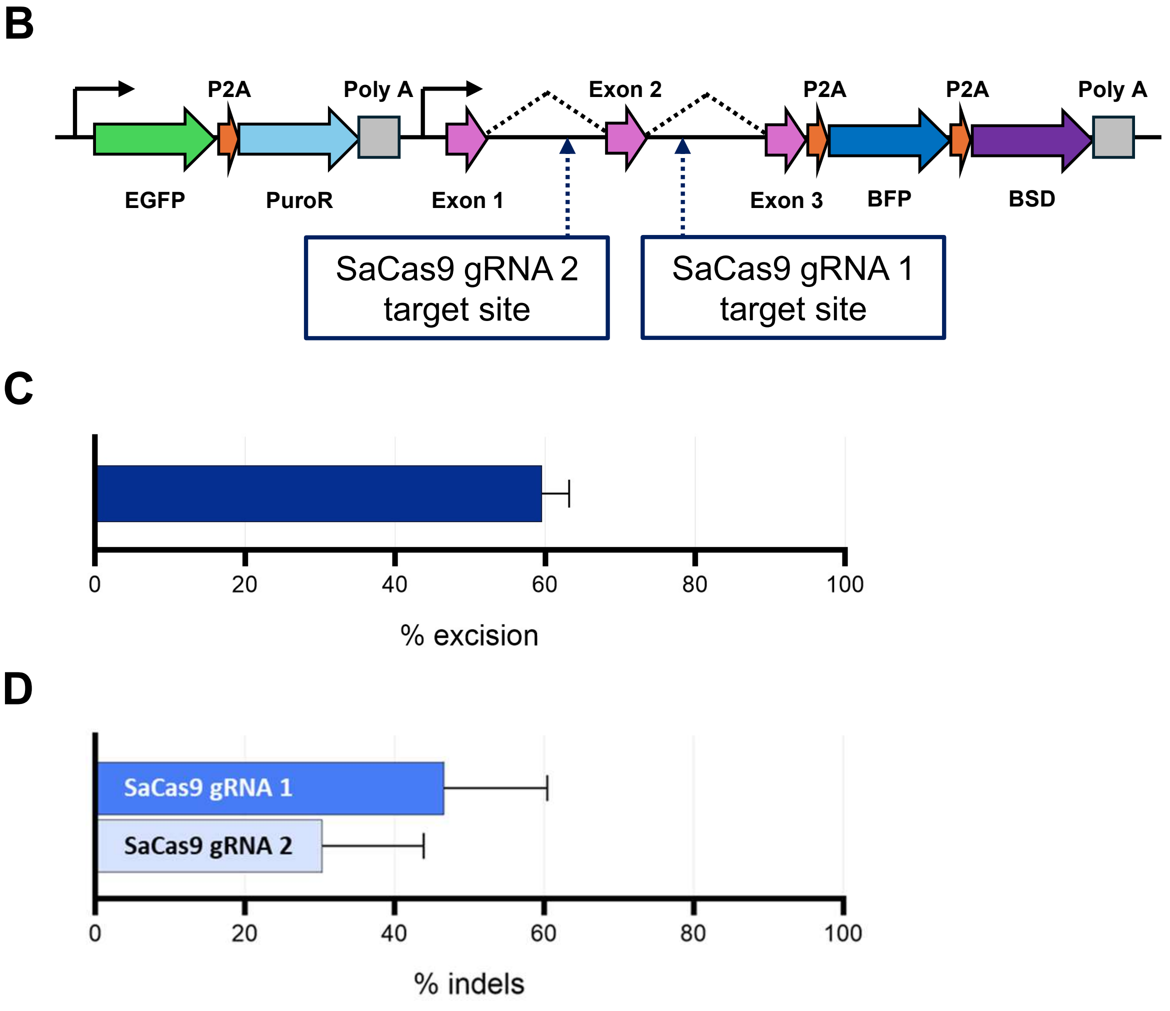


**Supplementary Figure 1. Characterization of paired guide RNAs selected to cut HBV genomic sequences.** (A) Conservation of the target sequences for the two HBV-targeting gRNAs, SaCas9 gRNA 1 and SaCas9 gRNA 2 were analyzed in silico using viral sequences sorted from HBVdb (https://hbvdb.lyon.inserm.fr/HBVdb/HBVdbIndex). The table shows the percentages of the viral target sites that contain no base-pair substitution and gap in the protospacer and PAM regions. (B) A schematic of the synthetic construct used to generate the HBV reporter cell line. The two HBV on-target sites separated by a synthetic exon with stop codons were inserted in the AAVS-1 locus (see Materials and Methods for details). (C) Excision of the intervening sequence between the two intended target sites was quantified by multiplex digital PCR in the HBV reporter cell line transfected with the SaCas9-encoding mRNA and a pair of the HBV-targeting gRNAs. (C) Indels at each of the target sites that were not lost by excisions or inversions were quantified by sequencing PCR-amplified fragments from the edited cells.

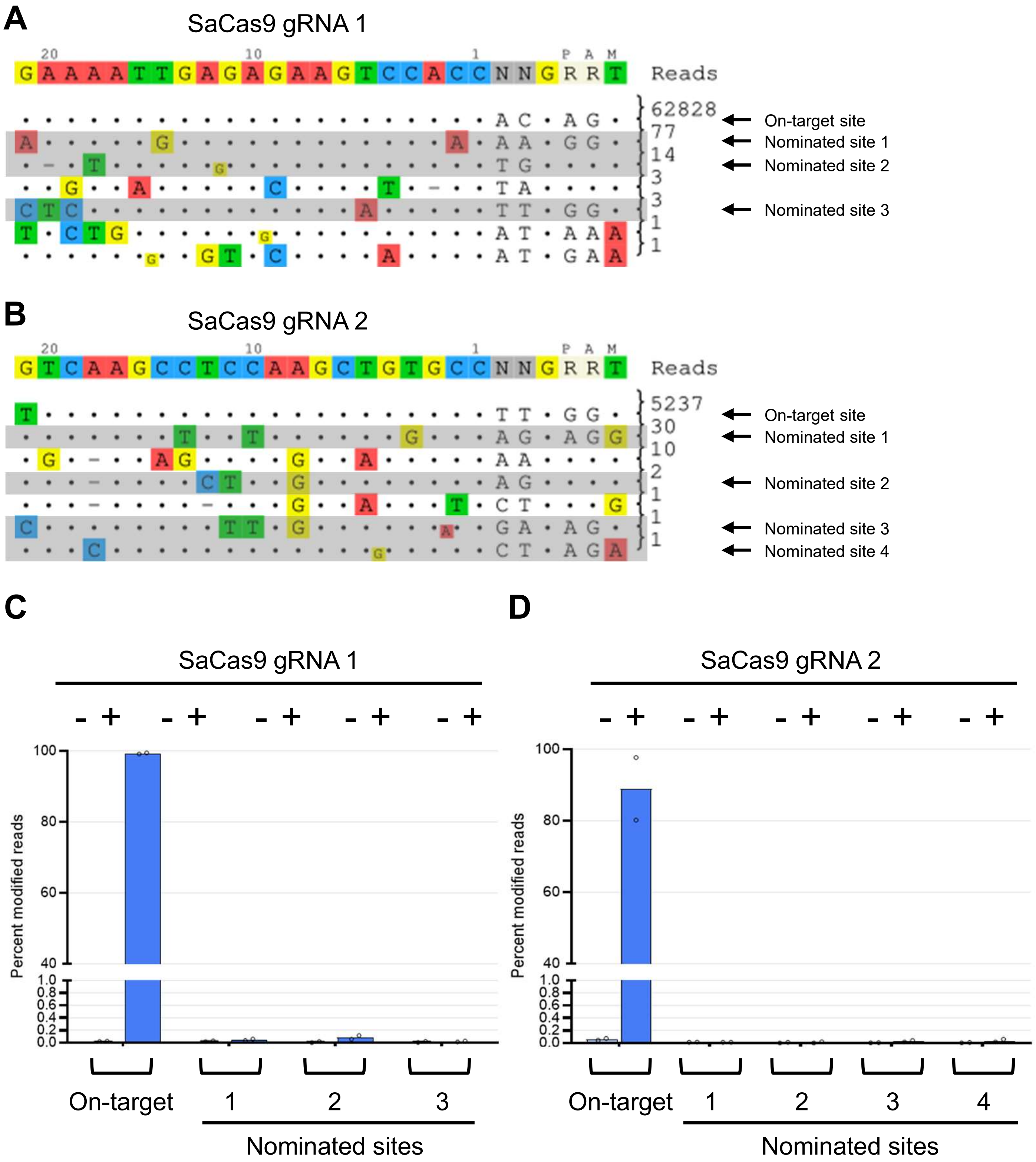


**Supplementary Figure 2. Sequence specificity of SaCas9 complexed with one of the paired HBV-targeting gRNAs.** (A,B) Human genomic sites nominated by GUIDE-seq for SaCas9 gRNA 1 and SaCas9 gRNA 2. The numbers of sequence reads detected at the on-target sites on the chromosomally integrated reporter construct are shown in the first row under each target sequence. Nominated sites for further off-target analysis are highlighted in gray and indicated by arrows. (C, D) Targeted amplicon sequencing to quantify indels at the intended target sites and nominated sites.

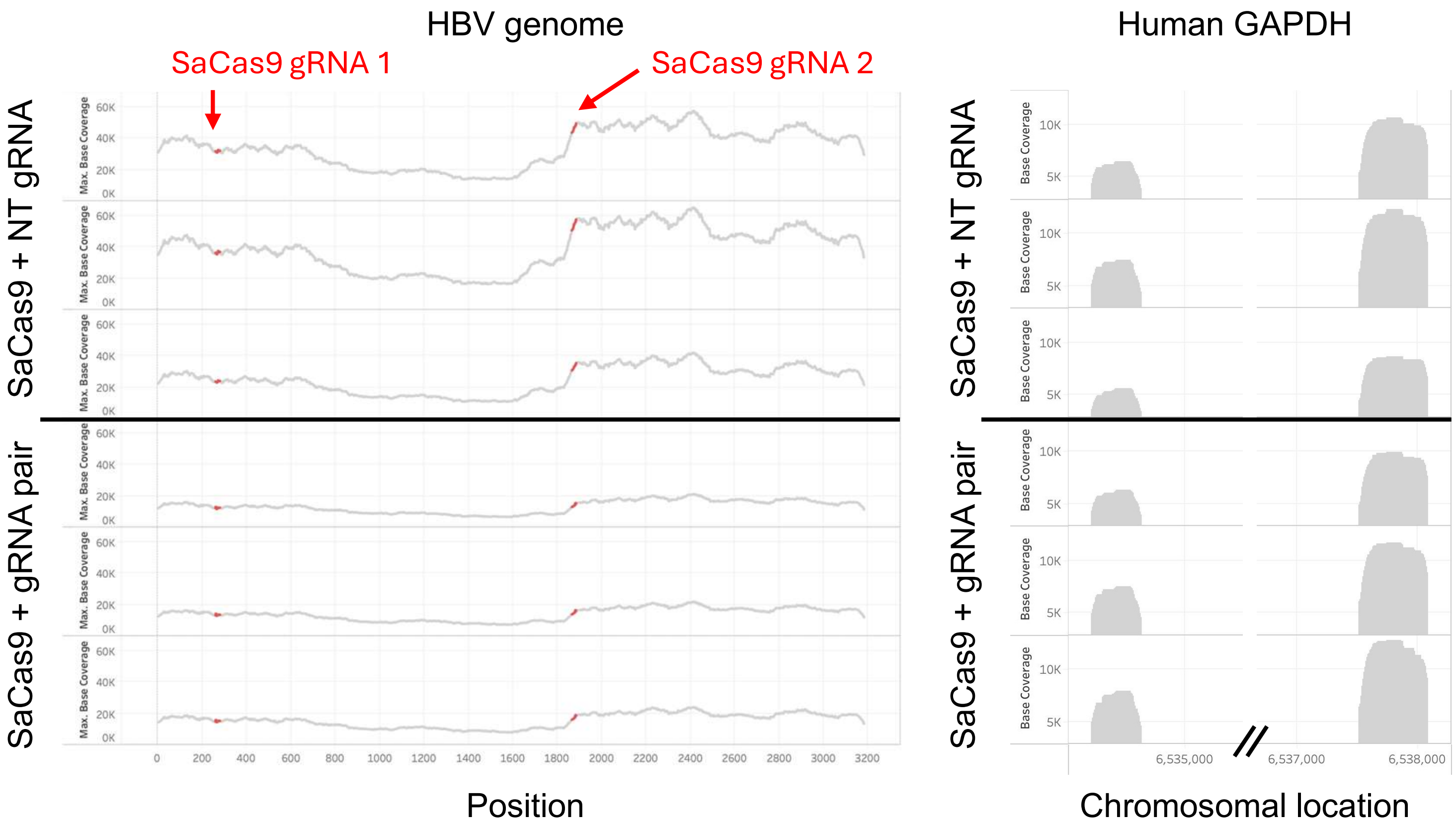


**Supplementary Figure 3. Sequence depth of HBV DNA extracted from HBV-infected PHHs by hybridization capture sequencing.** In addition to HBV DNA, two separate chromosomal regions within the human GAPDH gene locus were also targeted to obtain sequence reads that serve as the internal control. The HBV on-target sites are colored red.

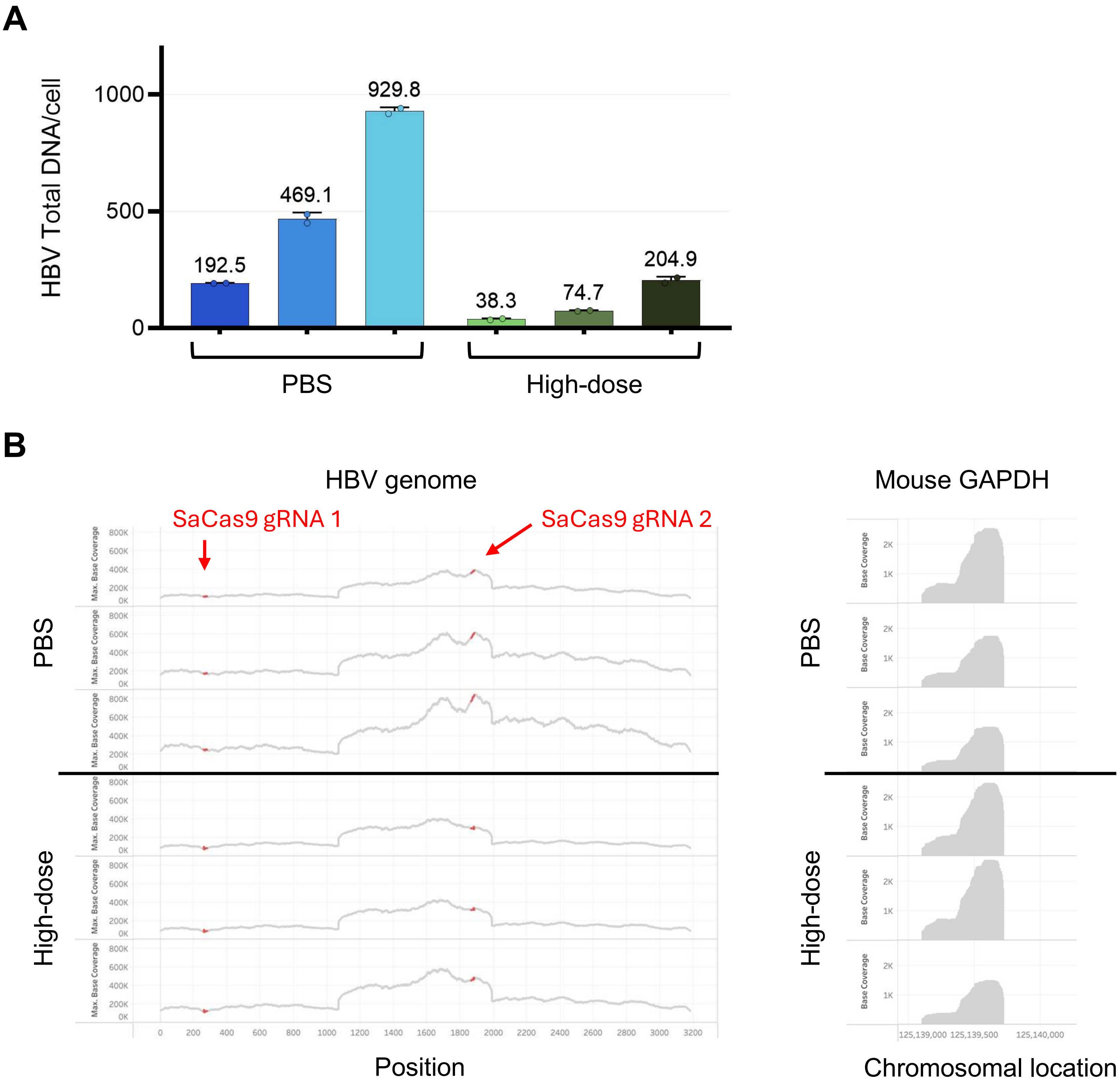


**Supplementary Figure 4. Copy number and sequence depth of HBV DNA extracted from liver of the AAV-HBV mice by hybridization capture sequencing.** (A) HBV total DNA in the samples subjected to hybridization capture sequencing was quantified by digital PCR. (B) In addition to HBV DNA, two separate chromosomal regions within the human GAPDH gene locus were also targeted to obtain sequence reads that serve as the internal control. The HBV on-target sites are colored red.

**A**

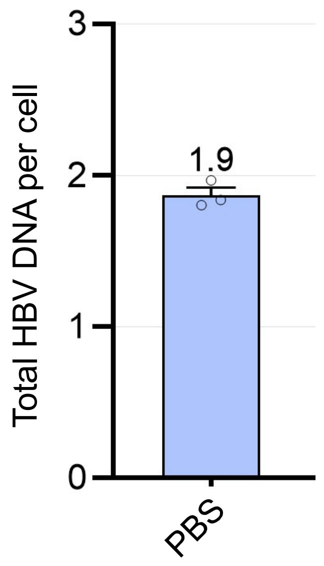


**B**

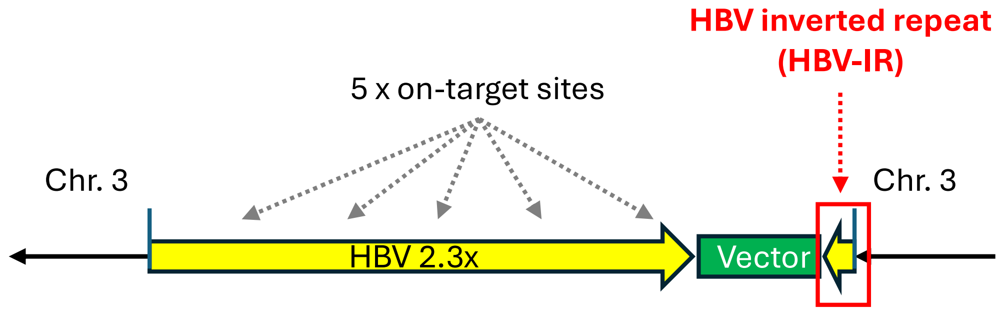


**Supplementary Figure 5. Structure of the chromosomally integrated transgenic HBV (Tg-HBV) sequence.** (A) Total HBV DNA copies were quantified by digital PCR using genomic DNA extracted from stomach tissue of one of the PBS-treated (control) Tg-HBV mice, where no viral replication was expected. The template DNA was digested with the AvaI restriction endonuclease that cuts a single site per HBV genome. (B) Long-read hybridization capture sequencing indicated that 2.3 x tandem repeats of the genotype-A (GenBank ID: AF305422.1) was inserted in the reverse orientation at 124,418,553-124,418,556 of the mouse chromosome 3. This region contained total 5 of the HBV on-target sites and was joined to the partial cloning vector sequence and the 240-bp HBV inverted repeat (HBV-IR; positions 2631-2870 of the nucleotide sequence with GenBank ID of AF305422.1).
